# Structural basis for HCMV Pentamer recognition by antibodies and neuropilin 2

**DOI:** 10.1101/2021.03.25.436804

**Authors:** Daniel Wrapp, Xiaohua Ye, Zhiqiang Ku, Hang Su, Harrison G. Jones, Nianshuang Wang, Akaash K. Mishra, Daniel C. Freed, Fengsheng Li, Aimin Tang, Leike Li, Dabbu Kumar Jaijyan, Hua Zhu, Dai Wang, Tong-Ming Fu, Ningyan Zhang, Zhiqiang An, Jason S. McLellan

## Abstract

Human cytomegalovirus (HCMV) encodes for multiple surface glycoproteins and glycoprotein complexes^1, 2^. One of these complexes, the HCMV Pentamer (gH, gL, UL128, UL130 and UL131), mediates tropism to both epithelial and endothelial cells by interacting with the cell surface receptor neuropilin 2 (NRP2)^3, 4^. Despite the critical nature of this interaction, the molecular determinants that govern NRP2 recognition remain unclear. Here we describe the cryo-EM structure of NRP2 bound to the HCMV Pentamer. The high-affinity interaction between these proteins is calcium-dependent and differs from the canonical C-terminal arginine (CendR) binding that NRP2 typically utilizes^5, 6^. The interaction is primarily mediated by NRP2 domains a2 and b2, which interact with UL128 and UL131. We also determine the structures of four human-derived neutralizing antibodies in complex with the HCMV Pentamer to define susceptible epitopes. The two most potent antibodies recognize a novel epitope yet do not compete with NRP2 binding. Collectively, these findings provide a structural basis for HCMV tropism and antibody-mediated neutralization, and serve as a guide for the development of HCMV treatments and vaccines.

## MAIN TEXT

Human cytomegalovirus (HCMV) is a ubiquitous pathogen and congenital infection can cause debilitating and permanent birth defects^7–9^. Despite the severity of these infections and the prevalence of this pathogen, there are currently no FDA-approved vaccines and therapeutic options are limited^10–12^. HCMV is an enveloped, double-stranded DNA virus of the family *Herpesviridae*^13^. The surface of the viral membrane is decorated by several glycoprotein complexes that mediate viral entry and membrane fusion^14–16^. One of these complexes is the HCMV Trimer, composed of glycoproteins gH, gL, and gO^2, 17^. The HCMV Trimer mediates tropism for fibroblasts by binding platelet derived growth factor receptor alpha (PDGFRα)^18, 19^. The HCMV Trimer is also capable of mediating infection of a broader variety of cell types by interacting with transforming growth factor beta receptor 3 (TGFβR3)^4, 20^. The other critical tropism-determining complex is the HCMV Pentamer, which is composed of glycoproteins UL128, UL130, UL131, and the same gH and gL proteins that comprise the bulk of the HCMV Trimer^1, 3^. This elongated heteropentamer mediates tropism for endothelial and epithelial cells by binding to neuropilin 2 (NRP2) and triggering the viral fusion protein, gB, to facilitate viral entry into host cells^4, 15, 21–23^.

Neuropilins 1 and 2 are single-pass transmembrane proteins that are expressed on the surface of neuronal, epithelial, and endothelial cells^24, 25^. Under normal conditions, these proteins function as receptors and co-receptors that engage in numerous physiological processes, including angiogenesis and development of the nervous system^26, 27^. NRP2 is composed of two N-terminal CUB domains (a1 and a2), two F5/8 domains (b1 and b2), a MAM domain, a transmembrane domain, and a C-terminal PDZ domain that is thought to mediate cytoplasmic signaling in response to extracellular stimuli^28, 29^. Perhaps the most thoroughly characterized of these stimuli is vascular endothelial growth factor (VEGF)^30^. The crystal structure of these proteins in complex with one another has been determined, revealing that the b1 domain of NRP2 engages the C-terminal arginine of VEGF^6, 31^. Since this initial characterization, the NRP2 b1 domain has been shown to interact with other binding partners via the same mechanism^32^, prompting the moniker “CendR” to refer to this exposed C-terminal arginine motif^5^. Although it has been shown that soluble NRP2 is capable of inhibiting HCMV infection of epithelial cells^4^, the molecular determinants that mediate this interaction remain unclear, and several additional Pentamer receptors have been proposed^33, 34^. The most potently neutralizing HCMV-directed antibodies are elicited against the Pentamer, suggesting that it represents a susceptible target for the development of vaccines and immunotherapeutics^35–37^.

To investigate NRP2 and mAb binding, we initiated structural and biophysical studies. Based on previous crystallographic experiments that reported conserved calcium-coordinating loops in both the a1 and a2 domains of NRP2^38, 39^, we measured the affinity of recombinantly expressed NRP2 a1a2b1b2 for the soluble HCMV Pentamer ectodomain in both the presence and absence of calcium. We found that in the presence of 2 mM EDTA, no association between NRP2 and Pentamer could be detected. However, when the same experiment was performed in the presence of 2 mM CaCl_2_, the affinity of the interaction was determined to be 2.2 nM (Supplementary Fig. 1a-b). It is possible that a failure to add additional calcium is what necessitated the use of chemical cross-linkers during previous efforts to observe this complex by negative-stain electron microscopy^4^. The addition of 2 mM CaCl_2_ enabled us to form a stable ∼230 kDa complex that was suitable for cryo-EM screening. The addition of 0.1% amphipol A8-35 helped to prevent aggregation and allowed for the determination of a 4.0 Å resolution cryo-EM structure of the HCMV Pentamer bound by human NRP2 (Fig. 1a, Supplementary Figs. 2 and 3). Performing focused refinement on the NRP2-bound UL proteins yielded a 3.65 Å reconstruction that aided in model building and refinement.

**Figure 1:**
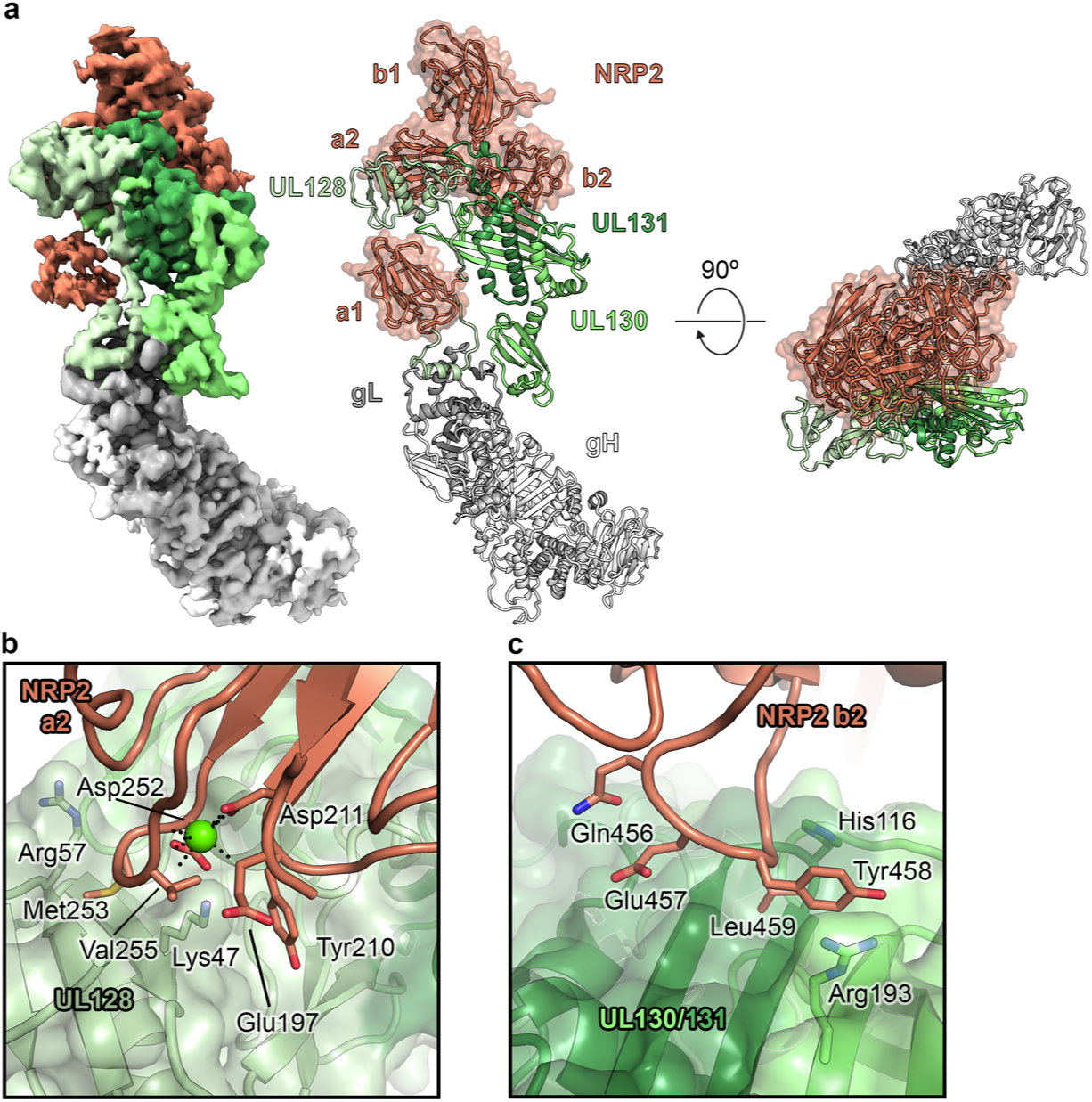
The cryo-EM structure of the HCMV Pentamer bound by NRP2. (**a**) Cryo-EM density is shown (*left*), with the Pentamer colored in shades of green, gray and white and NRP2 colored orange. The atomic model of this complex (right) is shown as ribbons, with the NRP2 also represented by a transparent molecular surface. (**b**) The interface between the NRP2 a2 domain and UL128. UL128 is depicted as a transparent, green molecular surface with ribbons underneath and NRP2 is shown as orange ribbons. Residues that are predicted to form critical contacts are shown as sticks. Oxygen, nitrogen, and sulfur atoms are colored red, blue, and yellow, respectively. The single calcium atom is shown as a bright green sphere, with black dotted lines depicting the interaction with conserved coordinating residues. (**c**) The interface between the NRP2 b2 domain and the HCMV Pentamer. ULs 130 and 131 are shown as a transparent, green molecular surface and NRP2 is shown as orange ribbons, with residues predicted to form critical contacts shown as sticks. Oxygen and nitrogen atoms are colored red and blue, respectively.

These reconstructions revealed an extensive binding interface, with contacts formed by NRP2 domains a1, a2 and b2 (Fig. 1). Notably, the calcium-coordinating loop of domain a2 (residues 251−258) forms a sizable portion of this binding interface, likely providing an explanation as to why high-affinity NRP2 binding could only be observed after the addition of 2 mM CaCl_2_ (Fig. 1a-b). Additional contacts are formed between the C-terminal beta strands of ULs 130 and 131 and a loop formed by residues 453-461 of the b2 domain of NRP2 (Fig. 1c). This mode of NRP2 binding differs from the canonical CendR motif binding that has been described previously for other NRP2-binding partners^40, 41^. The CendR binding mechanism involves the engagement of a C-terminal arginine residue by the b1 domain, whereas Pentamer is exclusively bound by the a1, a2, and b2 domains. Furthermore, none of the three UL proteins contain a positively charged C-terminal arginine that makes up the CendR motif. As expected, the majority of the binding interface from the Pentamer is composed of the tropism-determining UL proteins, particularly UL128 and UL131^1, 3^, which respectively contribute 437.5 Å^2^ and 208.4 Å^2^ of buried surface area to the interface. Whereas the NRP2 a2, b1, and b2 domains are clustered tightly together at the head of the Pentamer, the N-terminal a1 domain is tethered via a flexible linker that allows it to bind near the middle of the Pentamer, where the C-terminus of UL128 associates with gL. The local resolution for this portion of the reconstruction was relatively poor compared to the rest of the complex, suggesting either conformational flexibility in this region or a loose association of a1. To test the importance of the a1 domain, we expressed NRP2 with a 144-residue N-terminal truncation and observed that even in the absence of this flexibly tethered a1 domain, NRP2 a2b1b2 was capable of binding to the HCMV Pentamer with 7.9 nM affinity (Supplementary Fig. 1c), supporting our structural observations that the critical determinants of Pentamer binding are contained within NRP2 domains a2b1b2. Intriguingly, our cryo-EM data processing also revealed that a second, more poorly resolved copy of NRP2 could be observed binding near the C-terminal arginine of gL via the b1 domain (Supplementary Figs. 2 and 4). Although this second NRP2 appears to exhibit the canonical CendR binding, it could only be observed in ∼40% of particles. Furthermore, its binding to the gL protein rather than the tropism-determining UL proteins suggests that this second copy of NRP2 is likely an artifact of the high concentrations of NRP2 that were used to form a stable complex. Overall, the conformation of the receptor-bound Pentamer ectodomain does not drastically differ from that of the unbound Pentamer^42^ (Supplementary Fig. 5), suggesting that rather than undergoing substantial conformational rearrangements, this complex acts as a tether to connect HCMV virions to the surface of epithelial and endothelial cells until the viral fusogen gB fuses the viral and cellular membranes.

Previous efforts to characterize the humoral immune response to asymptomatic HCMV infection yielded an extensive panel of neutralizing antibodies directed against gB, the HCMV Trimer, and the HCMV Pentamer^35, 43, 44^. To learn more about the mechanisms of neutralization of high-affinity, Pentamer-directed antibodies, we determined cryo-EM structures of four naturally elicited human antibodies in complex with the Pentamer (Fig. 2, Supplementary Figs. 3, 6, 7, and 8). The flexibility and elongated shape of the Pentamer necessitated focused refinements of the Fabs along with the domains making up their respective epitopes. Model building was facilitated by high-resolution crystal structures of unbound Fabs (1-103: 1.9 Å, 1-32: 2.1 Å, 2-18: 2.8 Å, 2-25: 2.5 Å), which were then used as reference restraints and lightly refined as a part of the complex (Supplementary Tables 1 and 2). Three of these antibodies (1-103, 2-18, and 2-25) bound solely to the UL proteins at the head of the Pentamer, whereas the fourth (1-32) bound to gL, near the junction between the UL proteins and the conserved gH/gL scaffold (Fig. 2a). The Fab 1-103 epitope is solely composed of residues from the membrane-distal tip of UL128, sometimes referred to as Site 1 of immunogenic region (IR) 1^4, 35^. The epitopes of Fabs 2-18 and 2-25 overlap substantially, with both Fabs binding to the junction between ULs 128 and 131. This junction where UL128 meets UL131 does not fit into one of the preexisting antigenic sites, but rather overlaps with both Site 2 and Site 5 of IR1^4, 35^. The Fab 1-32 epitope spans the interface between gH and gL, slightly below Site 4/6 in IR2^4, 35^ (Fig. 2b). This epitope is consistent with previous observations^35^ that 1-32 is the only one of the four antibodies evaluated that was capable of binding to both the fully assembled HCMV Pentamer and disulfide-linked homodimers of the gH/gL heterodimer. Despite the ability to recognize the gH/gL heterodimer that serves as the scaffold for assembly of both the HCMV Pentamer and Trimer, 1-32 was only capable of neutralizing HCMV infection in epithelial cells^35^.

**Figure 2:**
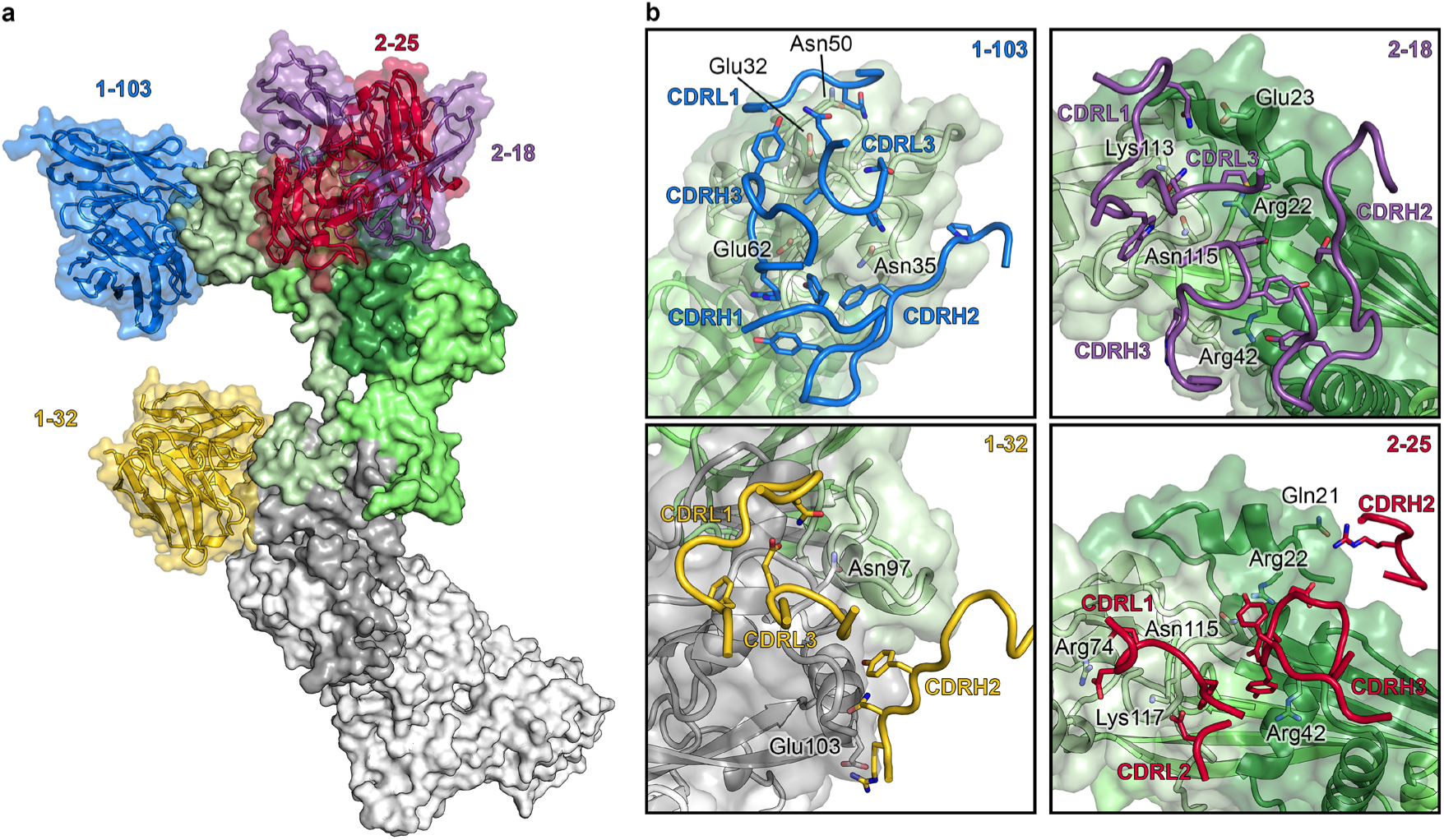
Composite of cryo-EM structures of Pentamer bound by four neutralizing human antibodies. (**a**) The atomic models of two cryo-EM structures of antibodies bound to the HCMV Pentamer are superimposed based on the position of the UL proteins. The Pentamer is shown as a molecular surface, colored according to Fig. 1 and Fabs are shown as ribbons surrounded by a transparent molecular surface. Fab 1-103 is colored blue, Fab 1-32 is colored gold, Fab 2-18 is colored purple and Fab 2-25 is colored red. (**b**) CDRs from each Fab are shown as ribbons and the Pentamer is shown as a transparent, molecular surface with ribbons underneath. Predicted critical contact residues are shown as sticks. Fab 1-103 (*top left*) is colored blue, Fab 1-32 (*bottom left*) is colored gold, Fab 2-18 (*top right*) is colored purple and Fab 2-25 (*bottom right*) is colored red. Oxygen, nitrogen and sulfur atoms are colored red, blue and yellow respectively.

By analyzing the structures of these immunocomplexes in conjunction with the structure of Pentamer bound by NRP2, it becomes possible to delineate more clearly the molecular basis for neutralization (Fig. 3). The CDR H3 and CDR L1 of Fab 1-103 compete with NRP2 by binding to the same portion of Pentamer that is engaged by the calcium-coordinating loop (residues 251 to 258) of NRP2 domain a2. By binding to the junction between gL and the UL head of Pentamer, Fab 1-32 occupies the same space as several of the loops of the a1 domain of NRP2 (residues 45−48; 72−77; 106−110) via its CDR L1 (Fig. 3b). Consistent with the relatively poor density for domain a1, we found that when bound by 1-32, Pentamer was still capable of interacting with NRP2, albeit with diminished affinity (133 nM) (Supplementary Fig. 9), further supporting our findings that the a1 domain is not strictly required for binding to Pentamer. This observation is also consistent with the relatively weak neutralization capacity of 1-32 compared to 1-103, which competes for the a2b1b2 interface.

**Figure 3:**
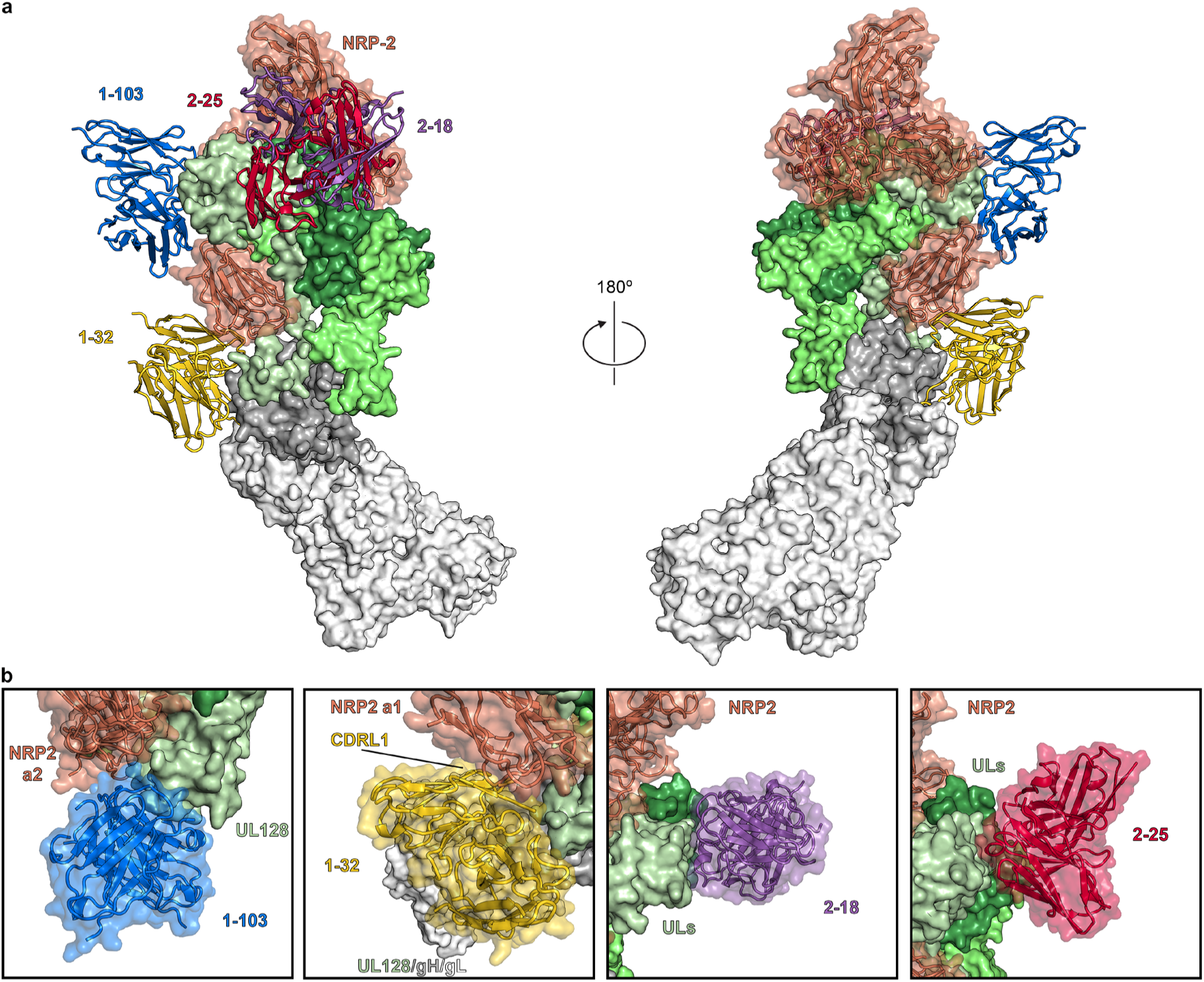
Pentamer-directed antibodies can neutralize HCMV via multiple mechanisms. (**a**) Cryo-EM structures of NRP2-bound Pentamer and Fab-bound Pentamer are superimposed based on the position of the UL proteins. Pentamer is shown as a molecular surface colored according to Fig. 1, Fabs are shown as ribbons, colored according to Fig. 2, and NRP2 is shown as orange ribbons surrounded by a transparent molecular surface. (**b**) Close-up views of each Fab-bound Pentamer are superimposed upon the NRP2-bound Pentamer. Both Fabs and NRP2 are shown as ribbons surrounded by a transparent molecular surface, while the Pentamer is shown as a solid molecular surface. NRP2 is colored orange, 1-103 is colored blue, 1-32 is colored gold, 2-18 is colored purple and 2-25 is colored red.

Intriguingly, the two most potently neutralizing mAbs that we examined, 2-18 and 2-25 (Fig. 4a), do not appear to compete with NRP2 a1a2b1b2 based on our structural analysis. Furthermore, although it is possible that 2-18 and 2-25 compete with the C-terminal MAM domain, this seems unlikely based on the position of the C-terminus of b2^45^. These mAbs are both directed against an epitope at the junction between ULs 128 and 131, and although this epitope is directly adjacent to the NRP2 binding interface, the binding angles of 2-18 and 2-25 result in these two Fabs being directed away from the β-sheet-rich face of the Pentamer that is responsible for engaging NRP2. A biolayer interferometry-based competition assay confirmed our structural observation that Fabs 2-18 and 2-25 do not disrupt the interaction between Pentamer and NRP2 a1a2b1b2 (Fig. 4b). These two Fabs are still capable of neutralizing HCMV *in vitro* even when administered up to 30 minutes after allowing viruses to adhere to human epithelial cells, although we observed a significant decrease in the neutralization potency of 2-25 Fab, as compared to 2-25 IgG (Figs. 4a, 4c). This broad and potent neutralization of multiple HCMV strains (Fig. 4d), which occurs without disrupting the interaction between NRP2 a1a2b1b2 and the Pentamer, suggests that 2-18 and 2-25 neutralize via a distinct mechanism from 1-103 and 1-32 (Supplementary Fig. 10). Whether 2-18 and 2-25 prevent the association of an unidentified secondary receptor or prevent some conformational change that is required to trigger gB-induced membrane fusion remains unclear and requires additional investigation.

**Figure 4:**
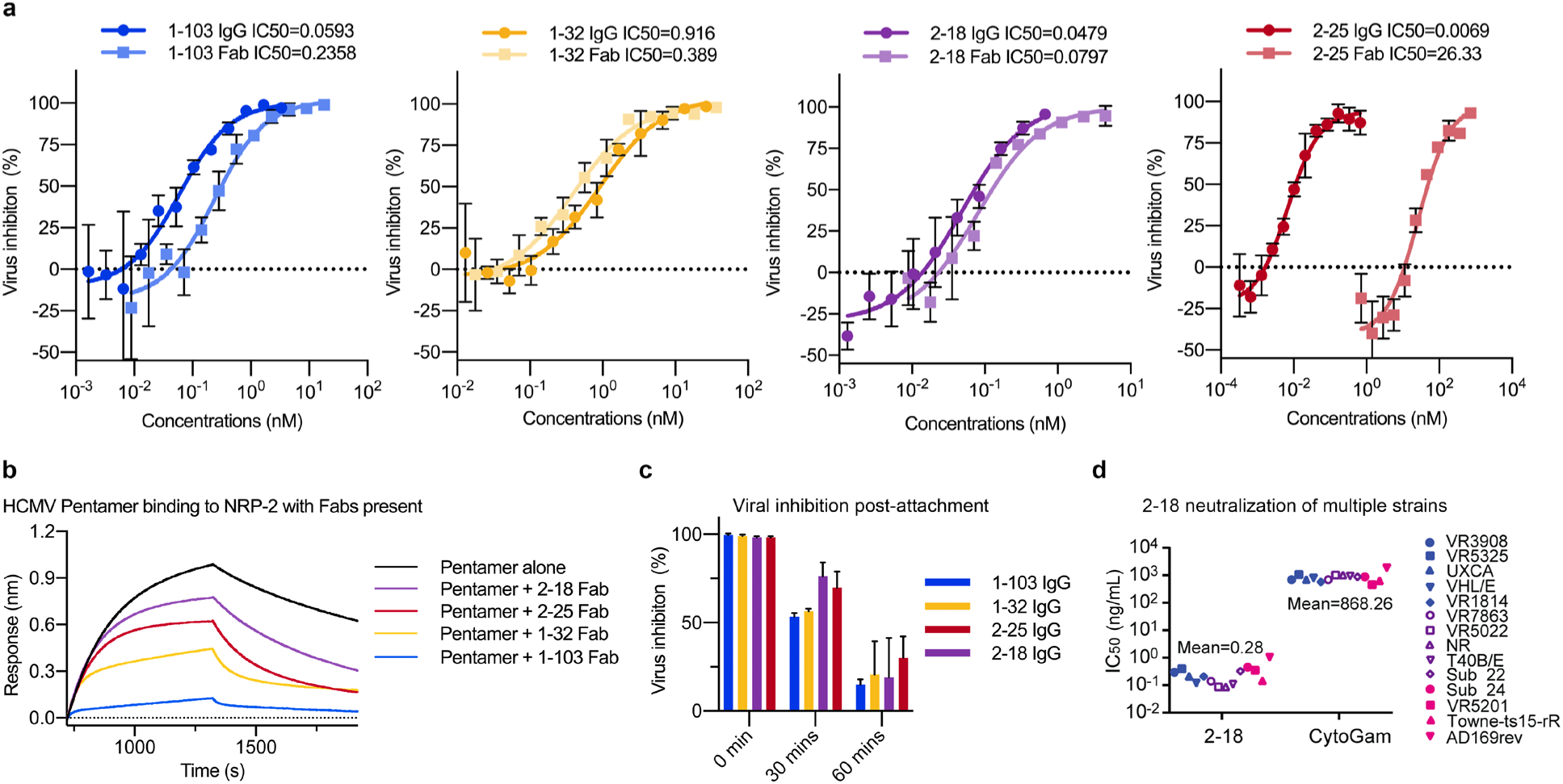
Antibodies 2-18 and 2-25 potently neutralize HCMV without disrupting NRP2 binding. (**a**) Neutralization curves are shown for each mAb based on inhibition of AD169rev-GFP infection in ARPE-19 cells. Inhibitory curves for both IgG and Fab are shown, with IgG shown in darker colors. (**b**) Sensorgrams from a BLI-based competition experiment are displayed. NRP2 was immobilized to a BLI sensor and dipped into Pentamer alone or Pentamer incubated with a molar excess of indicated Fab. (**c**) IgG neutralization of HCMV post-attachment to epithelial cells. AD169rev-GFP virions were allowed to adhere to ARPE-19 cells and saturating concentrations of IgG were added after a variety of different incubation periods. Viral inhibition is plotted for each IgG after either 0 mins incubation, 30 mins incubation, or 60 mins incubation. (**d**) Neutralization potency of 2-18 IgG was evaluated against twelve clinical isolates and two laboratory-adapted HCMV strains in ARPE-19 cells. IC_50_ values were calculated by non-linear fit of the percentage of viral inhibition vs. concentration (ng/mL). The neutralization results of mAbs 1-103, 1-32 and 2-25 against the same panel of HCMV strains have been reported previously^35^.

Collectively, these data provide a molecular basis of HCMV tropism for both epithelial and endothelial cells. Due to the importance of these cell types during natural infection, this represents a critical advance in our understanding of how HCMV engages host cells at one of the earliest stages of infection^22, 46^. Similarly, the structure of the HCMV Trimer was recently reported in complex with PDGFRα and TGFβR3, two host cell receptors that both mediate tropism of fibroblasts^18, 20^. In an effort to explain how this interaction might lead to triggering of gB and viral fusion, the authors speculate that receptor engagement of the Trimer may induce a rigid-body rotation relative to the viral membrane that causes the attachment complex to destabilize prefusion gB^47^. Our findings agree with their observation that receptor binding does not induce any conformation changes, lending credence to the hypothesis that a rigid-body rotation of the receptor-binding complex may act as trigger to induce membrane fusion. However, the existence of neutralizing mAbs that do not disrupt NRP2 binding suggests that additional fusion triggers, perhaps in the form of secondary receptors^33, 34^ may exist, necessitating further investigation.

In addition to detailing the molecular determinants that mediate HCMV infection, these findings expand our understanding of how antibody-mediated neutralization of HCMV can be achieved. Previous structural work has delineated a series of antigenic sites covering the surface of the Pentamer^4, 35, 42, 48^, but in the absence of high-resolution information regarding the NRP2 binding interface, the mechanisms of neutralization for antibodies targeting these sites were unknown. Our data suggest that it may be possible to neutralize HCMV via multiple, distinct mechanisms simultaneously by administering a cocktail of Pentamer-directed antibodies^36, 49, 50^. By elucidating which epitopes on the surface of the Pentamer are susceptible to antibody-mediated neutralization, these findings will also help to guide future structure-based vaccine design efforts.

## ACKNOWLEDGEMENTS

We thank Emilie Shipman and Dr. John Ludes-Meyers for their assistance with cell transfection and protein production; Drs. Aguang Dai and Sasha Dickinson at the Sauer Structural Biology Laboratory for their assistance with microscope alignment and data collection; the 19-ID beamline staff at the Structural Biology Center at the Advanced Photon Source, Argonne National Laboratory; and Dr. Georgina Salazar for assistance with manuscript preparation. This study was funded in part by grants from Merck & Co., Inc. Kenilworth, NJ, USA, the Texas Emerging Technology Fund, and the Welch Foundation Grant No. AU-0042-20030616. The Sauer Structural Biology Laboratory is supported by the University of Texas College of Natural Sciences and by award RR160023 from the Cancer Prevention and Research Institute of Texas (CPRIT). Argonne National Laboratory is operated by UChicago Argonne, LLC, for the U.S. Department of Energy (DOE), Office of Biological and Environmental Research under Contract DE-AC02-06CH11357.

## AUTHOR CONTRIBUTIONS

D.W., X.Y., Z.A., and J.S.M. conceived of and designed experiments. D.W., X.Y., H.G.J., N.W., and A.K.M. produced and purified proteins. D.W., H.G.J., and A.K.M. performed crystallographic studies. D.W. performed BLI and SPR experiments. D.W. and J.S.M. collected and analyzed cryo-EM data. X.Y., Z.K., and L.L. established the neutralization assays and X.Y., H.S., D.C.F., and F.L. performed the assays. X.Y., Z.K., A.T., and D. Wang analyzed the neutralization data. D.K.J. and H.Z. prepared the HCMV AD169rev-GFP virus. Z.A., T-M.F., N.Z., and J.S.M. supervised experiments. D.W., X.Y., Z.A., and J.S.M. wrote the manuscript with input from all authors.

## METHODS

### Protein production and purification

Plasmids encoding the heavy and light chains of 1-103, 1-32, 2-18, 2-25 and 8I21 IgG with an HRV3C protease cleavage site engineered into the hinge between the CH1 and CH2 domains of the heavy chain were co-transfected into FreeStyle 293F cells using polyethylenimine. To produce the soluble ectodomain of the HCMV Pentamer (strain Towne), plasmids encoding residues 24-718 of gH with a C-terminal 6x HisTag, residues 31-278 of gL, residues 21-171 of UL128, residues 26-214 of UL130 and residues 19-129 of UL131, all with artificial signal sequences were simultaneously co-transfected at an equimolar ratio.

Similarly, plasmids encoding an artificial signal peptide, residues 23-595 of human NRP2 and a C-terminal HRV3C cleavage site with either an 8x HisTag and a TwinStrepTag or a monomeric IgG1 Fc tag and an 8x HisTag were transfected into FreeStyle 293F cells, as described above. An N-terminal truncation of NRP2 that encompassed residues 145-595 with an artificial signal sequence and a C-terminal HRV3C cleavage site with a monomeric IgG1 Fc tag and an 8x HisTag (NRP2 a2b1b2) was transfected using the same conditions. NRP2 and NRP2 a2b1b2 were purified from cell supernatants using either StrepTactin resin (IBA) or Protein A resin before being run over a Superdex 200 Increase column using a buffer composed of 2 mM Tris pH 8.0, 200 mM NaCl, 0.02% NaN_3_ and 2 mM CaCl_2_.

To form the complex of Pentamer + 1-103 + 1-32 + 2-25, purified 1-103 IgG was immobilized to Protein A resin and this 1-103 resin was then used to capture Pentamer from co-transfected cell supernatants. The 1-103 + Pentamer complex was then eluted by incubation with HRV3C protease and purified over a Superose 6 Increase column in 2 mM Tris pH 8.0, 200 mM NaCl and 0.02% NaN_3_. This complex was then passed over a column containing 2-25 IgG immobilized to Protein A resin. Again, the complex was eluted by incubation with HRV3C protease and a molar excess of 1-32 Fab was added before a final round of purification over a Superose 6 Increase column using the same buffer.

To form the Pentamer + 2-18 + 8I21 complex, purified 2-18 IgG was immobilized to Protein A resin and used to capture Pentamer from co-transfected cell supernatants. The 2-18 + Pentamer complex was then eluted by incubation with HRV3C protease and mixed with a molar excess of 8I21 Fab before being run over a Superose 6 Increase column in 2 mM Tris pH 8.0, 200 mM NaCl and 0.02% NaN_3_.

To form the Pentamer + NRP2 complex, purified Pentamer was mixed with a threefold molar excess of 8x His/TwinStrep-tagged NRP2 in a buffer composed of 2 mM Tris pH 8.0, 200 mM NaCl, 0.02% NaN_3_ and 2 mM CaCl_2_ and the two components were allowed to bind on ice for 1 hour. This mixture was then purified over a Superose 6 Increase column (Cytiva) using the same buffer.

### X-ray crystallographic studies

Purified IgGs 1-103, 1-32, 2-18 and 2-25 were incubated with 10% (wt/wt) His-tagged HRV3C protease on ice for 2 hours before being passed over Protein A and NiNTA resin to removed cleaved Fc and excess protease. The remaining Fab was purified by SEC using a Superdex 200 Increase column in 2 mM Tris pH 8.0, 200 mM NaCl and 0.02% NaN_3_ (1-132 and 2-18) or 2 mM Tris pH 8.0, 50 mM NaCl and 0.02% NaN_3_ (1-103 and 2-25).

1-103 Fab was concentrated to 15.00 mg/mL and used to prepare sitting-drop crystallization trays. Diffraction-quality crystals grew in a mother liquor composed of 2.1 M sodium formate, 25% PEG 3350, 0.1 M sodium acetate pH 4.5 and 0.1 M calcium chloride. 1-103 Fab crystals were cryoprotected using mother liquor supplemented with 20% glycerol before being plunge frozen into liquid nitrogen.

1-32 Fab was concentrated to 11.00 mg/mL and used to prepare hanging-drop crystallization trays. Diffraction-quality crystals were grown in 2.0 M ammonium sulfate, 0.2 M sodium chloride and 5% isopropanol. 1-32 Fab crystals were cryoprotected using mother liquor supplemented with 20% glycerol before being plunge frozen into liquid nitrogen.

2-18 Fab was concentrated to 12.00 mg/mL and used to prepare sitting-drop crystallization trays. Small crystalline needles, grown in 0.1 M HEPES pH 7.5 and 45% PEG 400 were used to perform microseed matrix screening, ultimately yielding diffraction-quality crystals in a mother liquor composed of 0.2 M ammonium acetate, 0.1 M sodium citrate tribasic dihydrate pH 5.6 and 30% PEG 4000. 2-18 Fab crystals were cryoprotected using mother liquor supplemented with 20% glycerol before being plunge frozen into liquid nitrogen.

2-25 Fab was concentrated to 15.4 mg/mL and used to prepare sitting-drop crystallization trays. Diffraction-quality crystals were grown in 30% PEG 4000, a mixture of 0.2 M divalent cations^51^ and 0.1 M BIS-TRIS pH 6.5. 2-25 Fab crystals were looped without cryoprotectant and directly plunge frozen into liquid nitrogen.

All diffraction data were collected at Argonne National Labs, Advanced Photon Source, SBC-19ID. Datasets were indexed in iMOSFLM^52^ and scaled in AIMLESS^53^. Molecular replacement solutions were determined using PhaserMR^54^ and models were iteratively built and refined using Coot^55^, PHENIX^56^ and ISOLDE^57^. Full crystallographic data collection and refinement statistics can be found in Supplementary Table 1. Crystallographic software packages were curated by SBGrid^58^.

### Cryo-EM sample preparation and data collection

Purified HCMV Pentamer + 2-18 + 8I21 complex was diluted to a concentration of 0.25 mg/mL in 2 mM Tris pH 8.0, 200 mM NaCl, 0.02% NaN_3_ and 0.01% amphipol A8-35. 8I21 Fab was added after initial attempts to visualize the Pentamer + 2-18 complex were hampered by a lack of distinguishable features (data not shown). 3 μL of the ternary complex was deposited on a CF-1.2/1.3 grid that was glow discharged at 25 mA for 1 minute using an Emitech K100X (Quorum Technologies). Excess liquid was blotted away for 6 seconds in a Vitrobot Mark IV (FEI) operating at 4° C and 100% humidity before being plunge frozen into liquid ethane. Data were collected on a Titan Krios (FEI) operating at 300 kV, equipped with a K3 direct electron detector (Gatan). Movies were collected using SerialEM^59^ at 22,500x magnification, corresponding to a pixel size of 1.047 Å.

Purified HCMV Pentamer + 1-103 + 1-32 + 2-25 complex was diluted to a concentration of 0.2 mg/mL in 2 mM Tris pH 8.0, 400 mM NaCl, 0.02% NaN_3_ and 0.01% amphipol A8-35. 3 μL of protein was deposited on a CF-1.2/1.3 grid that was plasma cleaned at 25 mA for 30 seconds using a Solarus plasma cleaner (Gatan). Excess liquid was blotted away for 6 seconds in a Vitrobot Mark IV (FEI) operating at 4° C and 100% humidity before being plunge frozen into liquid ethane. Data were collected on a Titan Krios (FEI) operating at 300 kV, equipped with a K2 direct electron detector (Gatan). Movies were collected using Leginon^60^ at 22,500x magnification, corresponding to a pixel size of 1.075 Å.

Purified HCMV Pentamer + NRP2 complex was diluted to a concentration of 0.4 mg/mL in 2 mM Tris pH 8.0, 200 mM NaCl, 2 mM CaCl_2_, 0.02% NaN_3_ and 0.01% amphipol A8-35. 3 μL of protein was deposited on an UltrAuFoil 1.2/1.3 grid that was plasma cleaned at 25 mA for 2 minutes using a Solarus plasma cleaner (Gatan). Excess liquid was blotted away for 3 seconds in a Vitrobot Mark IV (FEI) operating at 4° C and 100% humidity before being plunge frozen into liquid ethane. Data were collected on a Titan Krios (FEI) operating at 300 kV, equipped with a K3 direct electron detector (Gatan). Movies were collected using SerialEM^59^ at 22,500x magnification, corresponding to a pixel size of 1.073 Å.

### Cryo-EM data processing and model building

Motion correction, CTF-estimation and non-templated particle picking using BoxNet were performed in Warp^61^. Extracted particles were imported into cryoSPARC^62^ for 2D classification, *ab initio* 3D reconstruction calculation, 3D classification and non-uniform refinement^63^. Based on the flexibility of the interface between the gH/gL and UL proteins, particle subtraction and focused refinement were also performed in cryoSPARC. Final reconstructions were sharpened with DeepEMhancer^64^. A full description of the cryo-EM data processing workflows can be found in Supplementary Figs. 2, 6 and 7. Crystal structures were docked into cryo-EM density maps using Chimera^65^ before being refined in Coot^55^, PHENIX^56^ and ISOLDE^57^. A detailed description of the cryo-EM data processing workflow can be found in Supplementary Figs. 2, 6 and 7. Full cryo-EM data collection and refinement statistics can be found in Supplementary Table 2.

### Surface plasmon resonance (SPR)

Purified His-tagged Pentamer was immobilized to a single flow cell of a NiNTA sensor in a Biacore X100 (GE Healthcare) to a level of ∼800 response units (RUs) using HBS-P+ buffer adjusted to a pH of 8.0. Two-fold serial dilutions of Fabs 1-103, 1-32, 2-18 and 2-25 were injected over both flow cells to measure binding kinetics. The sensor was doubly regenerated using 350 mM EDTA and 100 mM NaOH in between cycles. Data were double reference-subtracted and fit to a 1:1 binding model using Biacore Evaluation Software (GE Healthcare).

### Biolayer Interferometry (BLI)

Purified monoFc-tagged NRP2 or NRP2 a2b1b2 was immobilized to anti-human capture (AHC) tips (ForteBio) in a buffer composed of 10 mM HEPES pH 8.0, 150 mM NaCl, 0.05% Tween 20, 1 mg/mL BSA and 2 mM CaCl_2_. Sensors were then dipped into wells containing purified HCMV Pentamer, ranging in concentration from 50 nM to 3.125 nM. Data were reference subtracted and processed using Octet Data Analysis software v10.0 (ForteBio) with a 1:1 binding model. To evaluate the impact of calcium on Pentamer binding, the same experiment was performed using monoFc-tagged NRP2 in a buffer composed of 10 mM HEPES pH 8.0, 150 mM NaCl, 0.05% Tween 20, 1 mg/mL BSA and 2 mM EDTA.

To evaluate competition between Fabs and NRP2, monoFc-tagged NRP2 was immobilized to AHC tips in a buffer composed of 10 mM HEPES pH 8.0, 150 mM NaCl, 0.05% Tween 20, 1 mg/mL BSA and 2 mM CaCl_2_. Sensors were then dipped into wells containing a mixture of purified HCMV Pentamer at a concentration of 50 nM and 100 nM Fab. Data were reference subtracted using Octet Data Analysis software v10.0.

To measure the binding kinetics of NRP2 to Pentamer in the presence of mAb 1-32, 1-32 IgG was immobilized to AHC sensors using a buffer composed of 10 mM HEPES pH 8.0, 150 mM NaCl, 0.05% Tween 20, 1 mg/mL BSA and 2 mM CaCl_2_. Tips with immobilized 1-32 were then dipped into wells containing 100 nM Pentamer. The 1-32-captured Pentamer was then dipped into wells containing untagged NRP2, ranging in concentration from 400 nM to 25 nM. Data were reference subtracted and processed using Octet Data Analysis software v10.0 with a 1:1 binding model.

### HCMV neutralization assay

All of the Pentamer-specific antibodies used for the purposes of neutralization and inhibition assays were produced as described previously^43^. A dengue virus specific human IgG1 antibody^66^ was used as isotype control. Fabs for neutralization and inhibition assays were generated by digesting IgG with papain (Sigma, P4762) and purifying as described previously^67^. A standard neutralization assay with the Towne-ts15-rR, AD169rev, and 12 clinical isolates as shown in Supplementary Fig. 10 were performed in ARPE-19 cells using an immunostaining method^68^. Neutralization assays in Fig. 4 were performed in ARPE-19 cells using AD169rev-GFP strain and virus infection was examined through GFP expression as described previously^69^. For the standard neutralization assay, 50 µL/well of AD169rev-GFP, generating about 100 GFP-positive cells was incubated with 50 µL/well of serial 2-fold diluted IgG or Fab (at indicated concentrations) at 37 °C for 30 min and then added to confluent ARPE-19 cells grown in a 96-well plate. Mock-infected cells and cells infected with virus-only served as controls. For the post-attachment assay, ARPE-19 cells grown in a 96-well plate were pre-cooled at 4 °C for 10 min. 50 µL/well of AD169rev-GFP was allowed to attach to cells for 1 h at 4 °C. After removing unattached virus through a single wash using cold media, the indicated IgG, diluted at concentrations of ∼200 times of corresponding IC_50_ was added after culturing AD169rev-GFP-attached cells for different lengths of time (0 min, 30 min, and 60 min) in a 37 °C incubator. The antibody-containing media was replaced with fresh media without antibody 2 h later. Mock-infected cells and cells infected with virus but not treated with antibodies served as controls. For all above assays, triplicate wells were determined for each condition and viral infection was examined at 48 h post-infection. A C.T.L. Immunospot analyzer was used to capture whole-well images of GFP expression and quantification of GFP-positive cells. The percentage of viral inhibition by the antibody and the IC50 of each antibody was calculated by non-linear fit of virus inhibition % vs. concentration (ng/mL) using GraphPad Prism® 5 software.

---

**Supplementary Figure 1:**
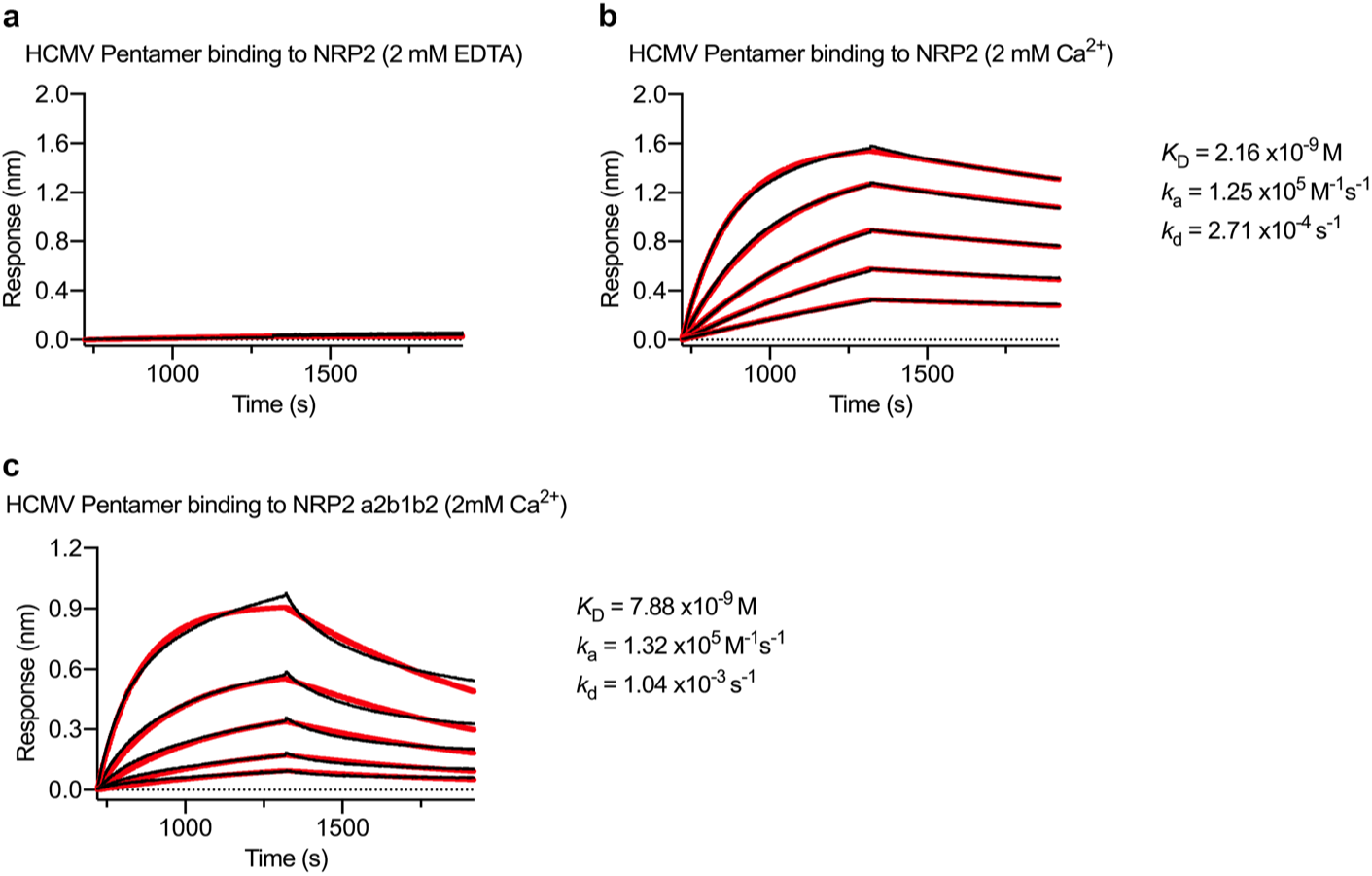
The interaction between NRP2 and Pentamer is calcium-dependent. (**a**) BLI sensorgram showing the absence of binding between Pentamer and NRP2 in the presence of 2 mM EDTA. (**b**) BLI sensorgram showing binding between Pentamer and NRP2 in the presence of 2 mM calcium. Data are shown as black lines and best fit of a 1:1 binding model is shown as red lines. (**c**) BLI sensorgram showing binding between Pentamer and NRP2 a2b1b2 in the presence of 2 mM calcium. Data are shown as black lines and best fit of a 1:1 binding model is shown as red lines.

**Supplementary Figure 2:**
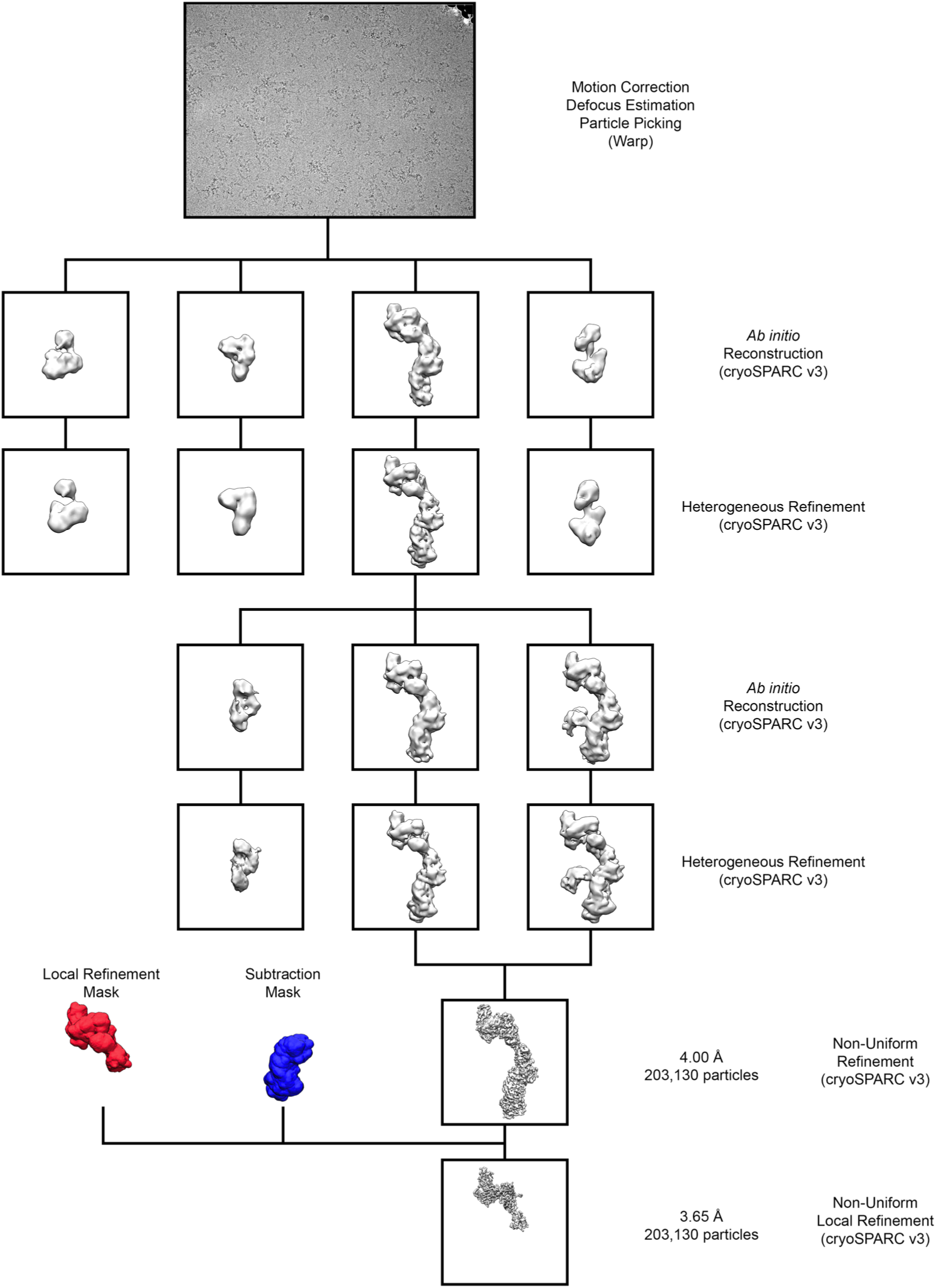
Pentamer + NRP2 cryo-EM data processing workflow.

**Supplementary Figure 3:**
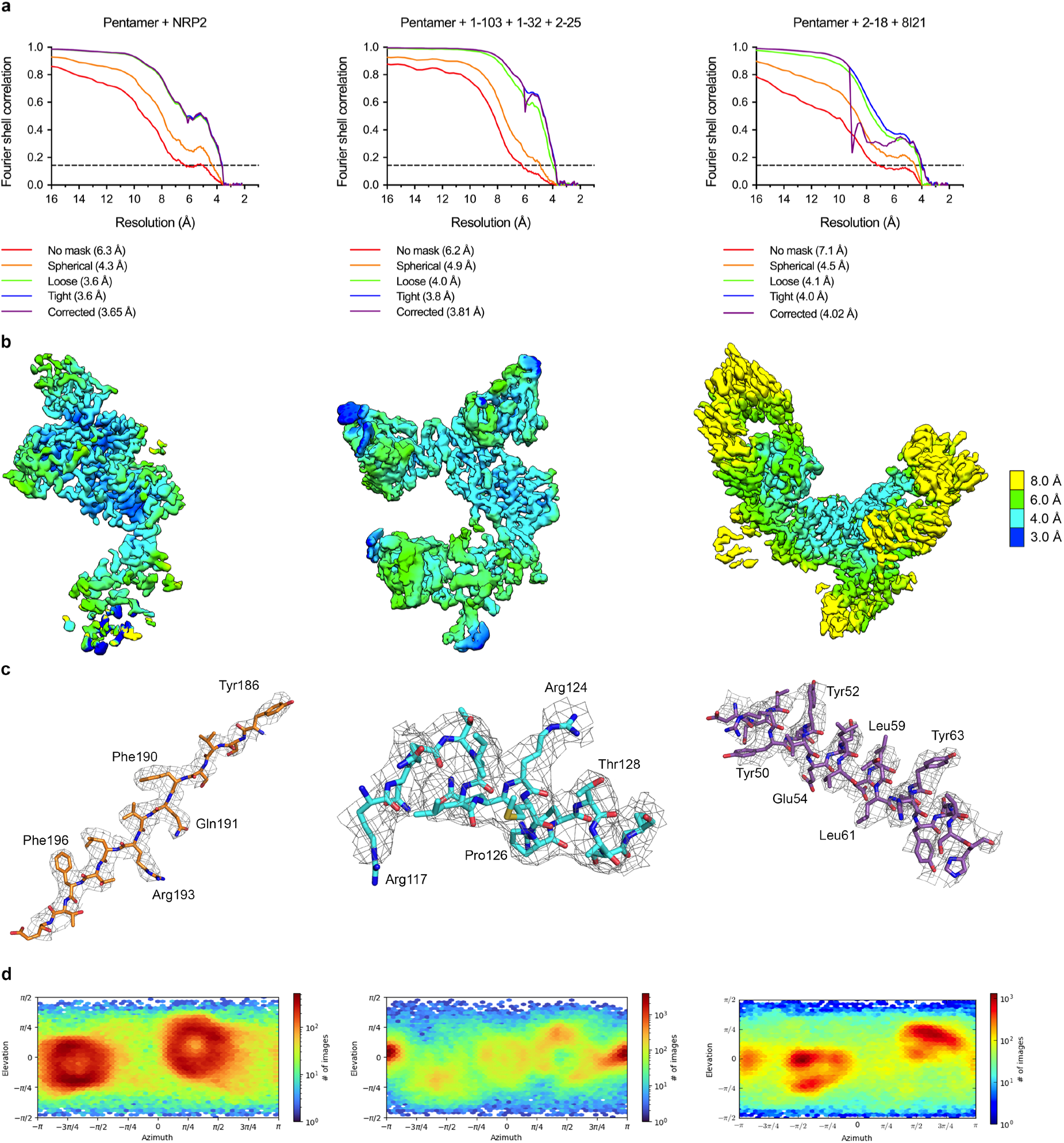
Cryo-EM structure validation. **(a)** FSC curves are shown for focused refinements of Pentamer bound by NRP2 (*left*), Pentamer bound by 1-103, 1-32 and 2-25 (*middle*) and Pentamer bound by 2-18 and 8I21 (*right*). **(b)** Cryo-EM maps from each focused refinement are shown, colored according to local resolution. **(c)** Portions of each cryo-EM map are shown, with the corresponding atomic models docked into the density. Residue numbering corresponds to UL130 (*left, middle*) and UL131 (*right*). (**d**) Viewing Direction Distribution charts from cryoSPARC are shown for each focused refinement.

**Supplementary Figure 4:**
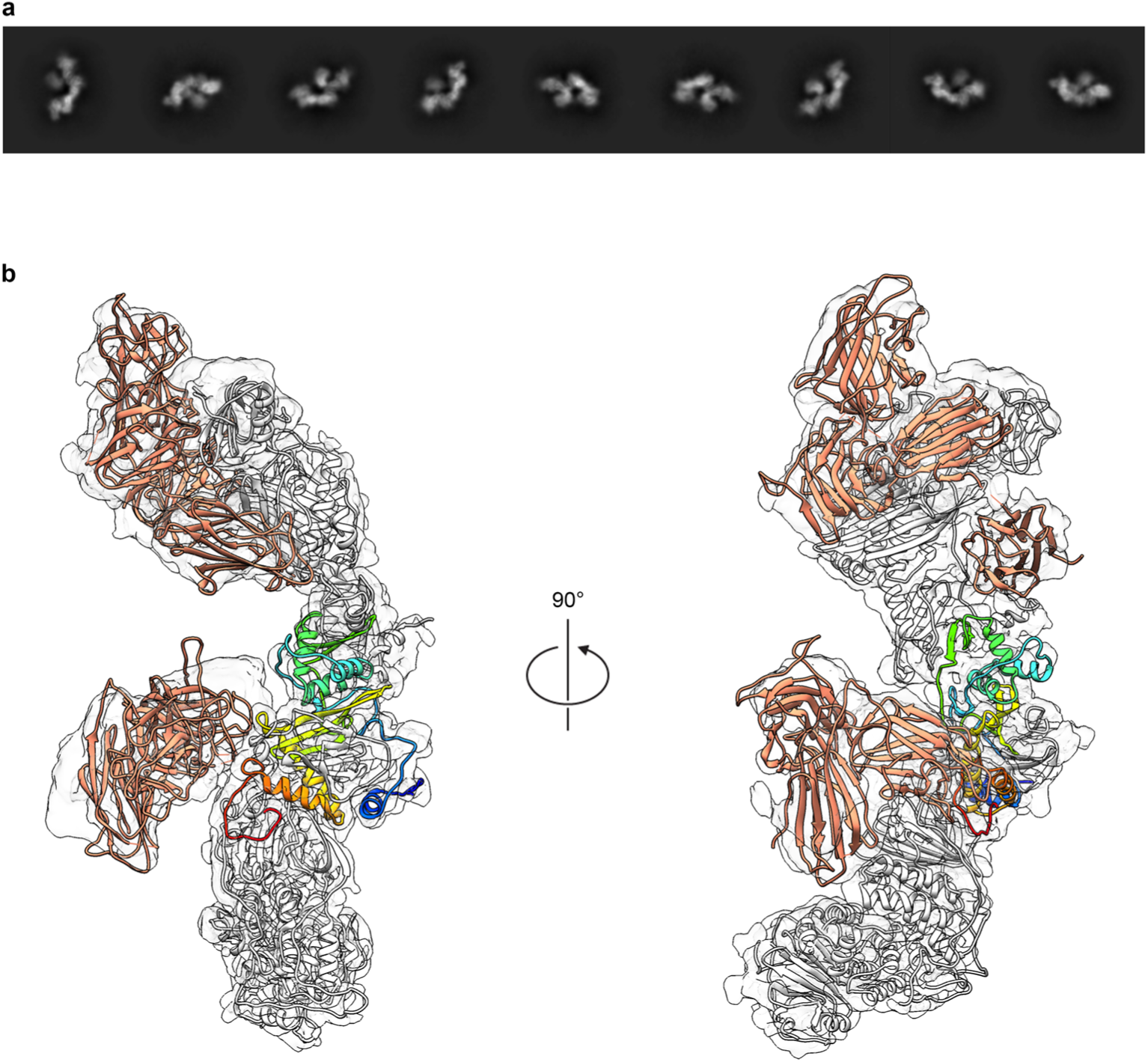
A subset of particles display a second copy of NRP2 bound to the C-terminus of gL. (**a**) Two-dimensional class averages of Pentamer bound by two copies of NRP2 are shown. (**b**) A ∼4.2 Å cryo-EM reconstruction of Pentamer bound by two copies of NRP2 is shown as a transparent surface. Atomic models of each component are docked in, shown as ribbons. Both copies of NRP2 are colored orange and Pentamer is colored white, except for gL, which is colored blue-to-red from the N-terminus to the C-terminus. The a1 domain from the gL-bound copy of NRP2 was excluded because it could not clearly be resolved in this reconstruction.

**Supplementary Figure 5:**
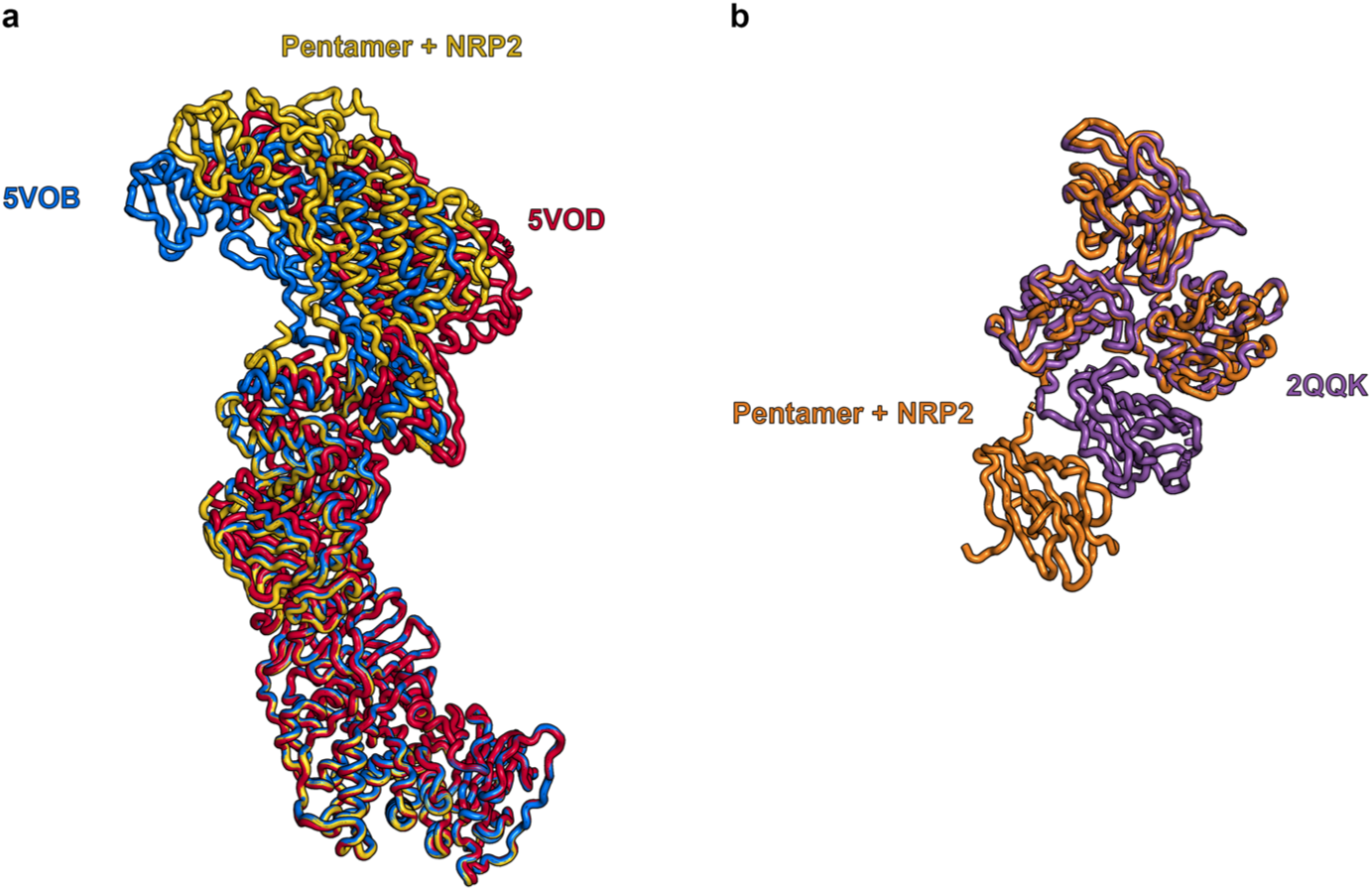
NRP2 binding does not alter the conformation of Pentamer. (**a**) Previously reported crystal structures of Pentamer^42^ (PDB IDs: 5V0B and 5V0D) are aligned to the cryo-EM structure of the NRP2-bound Pentamer, based on the position of gH. 5V0B is colored blue, 5V0D is colored red and NRP2-bound Pentamer is colored yellow. (**b**) A previously reported crystal structure of NRP2^38^ (PDB ID: 2QQK) is aligned to the cryo-EM structure of Pentamer-bound NRP2, based on the position of the b1 domain. 2QQK is colored purple and Pentamer-bound NRP2 is colored orange.

**Supplementary Figure 6:**
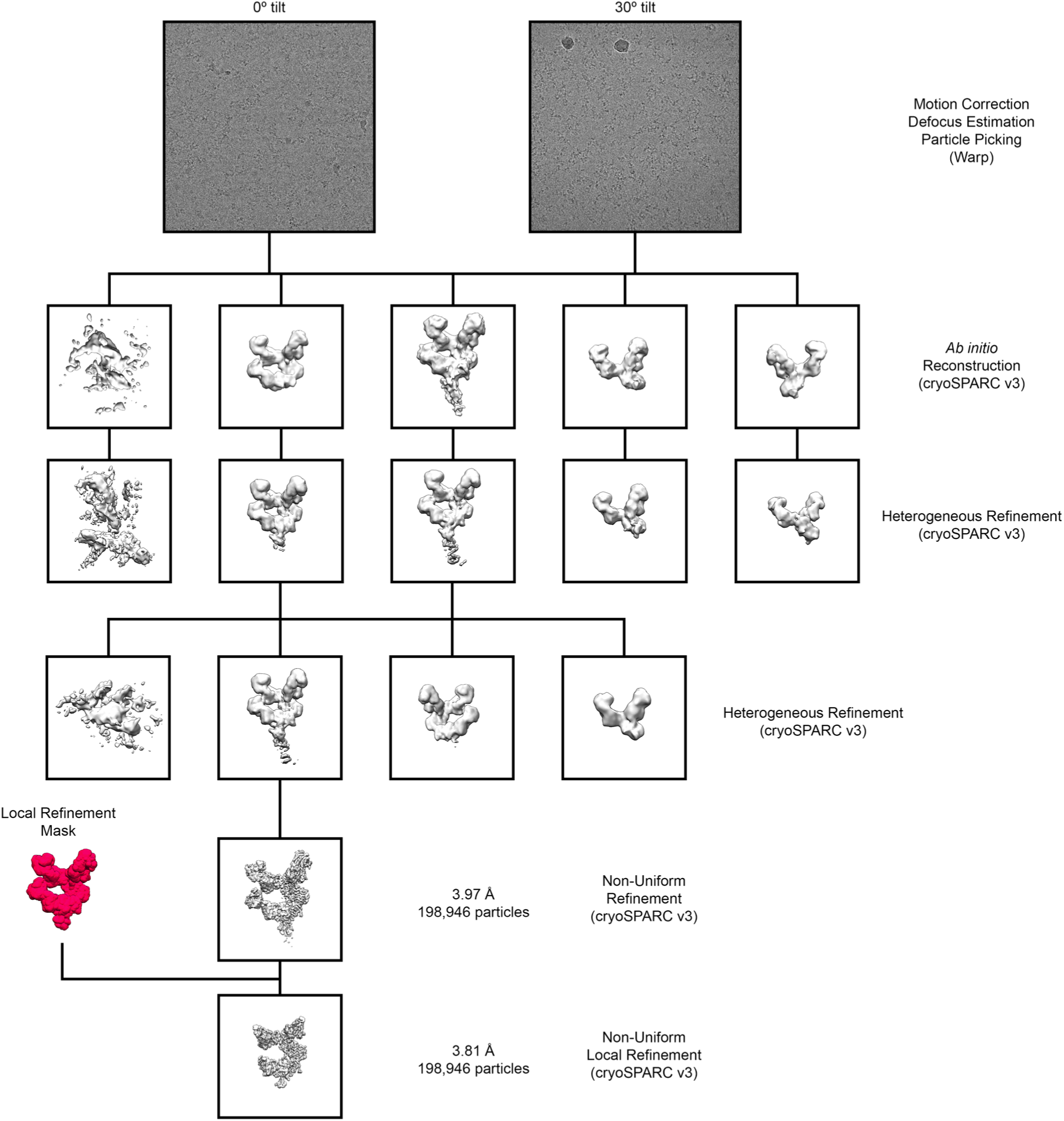
Pentamer + 1-103 + 1-32 + 2-25 cryo-EM data processing workflow.

**Supplementary Figure 7:**
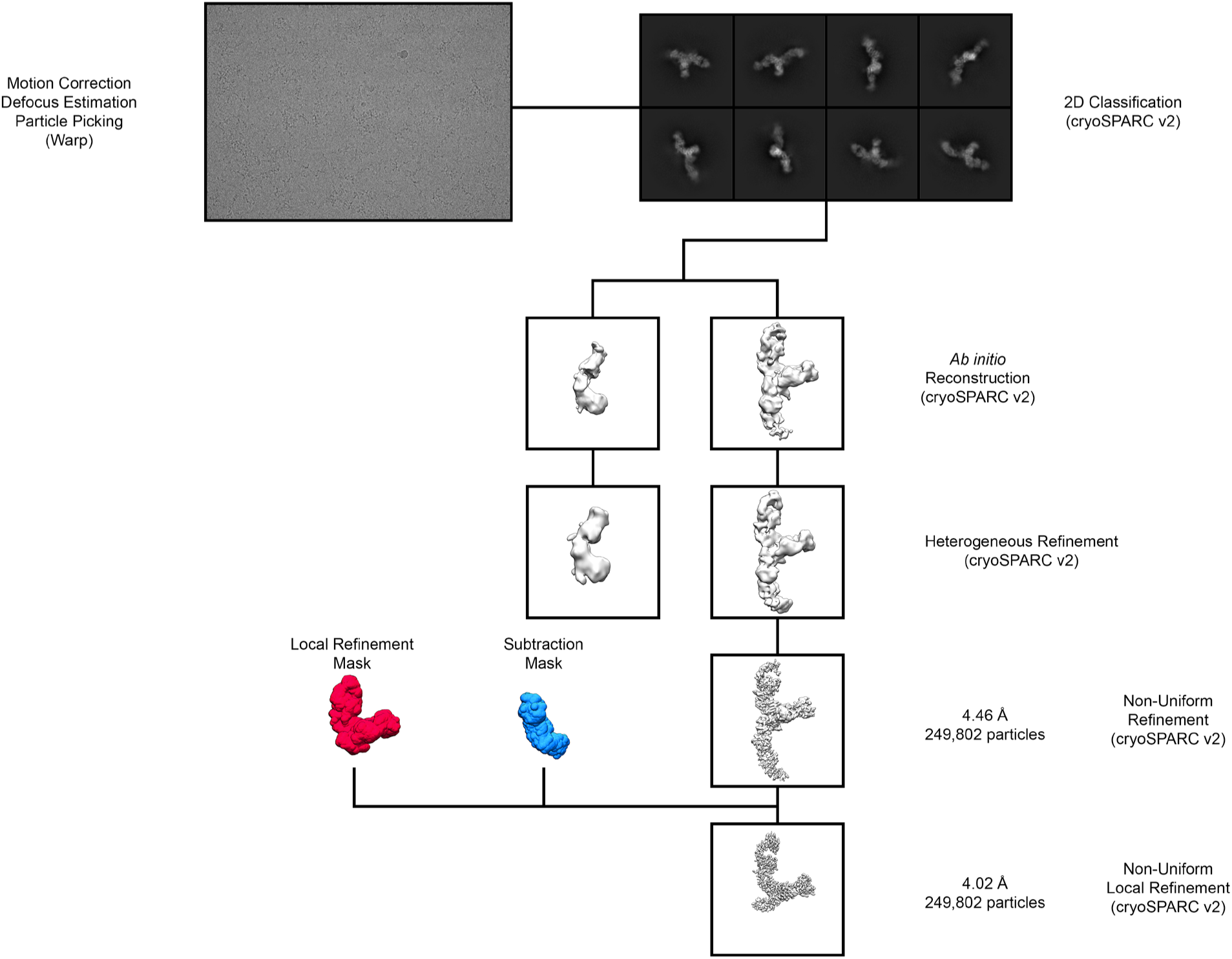
Pentamer + 2-18 + 8I21 cryo-EM data processing workflow.

**Supplementary Figure 8:**
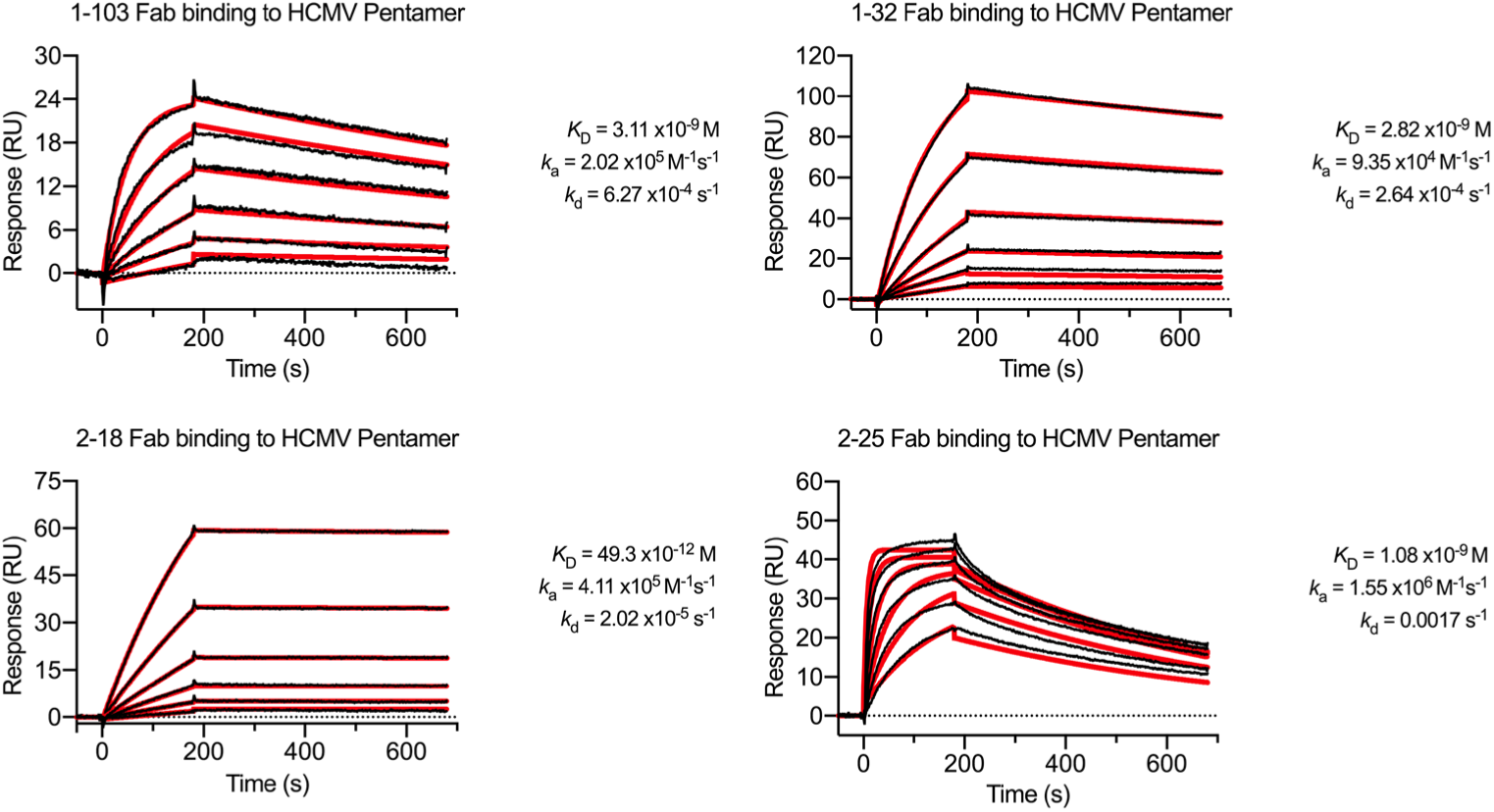
Binding kinetics of Pentamer-directed antibodies. SPR sensorgrams showing binding of each of the four neutralizing Fabs are displayed, with data shown as black lines and the best fit of a 1:1 binding model shown as red lines.

**Supplementary Figure 9:**
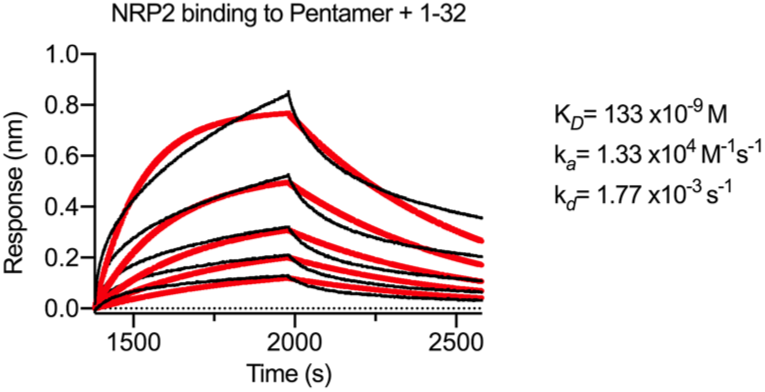
NRP2 binding to Pentamer is partially disrupted by the presence of 1-32. Sensorgrams are shown for an experiment in which 1-32 IgG was immobilized to a BLI sensor, then used to capture Pentamer before being dipped into NRP2. Data for the association and dissociation of NRP2 are shown as black lines and the lines of best fit of a 1:1 binding model are shown as red lines.

**Supplementary Figure 10:**
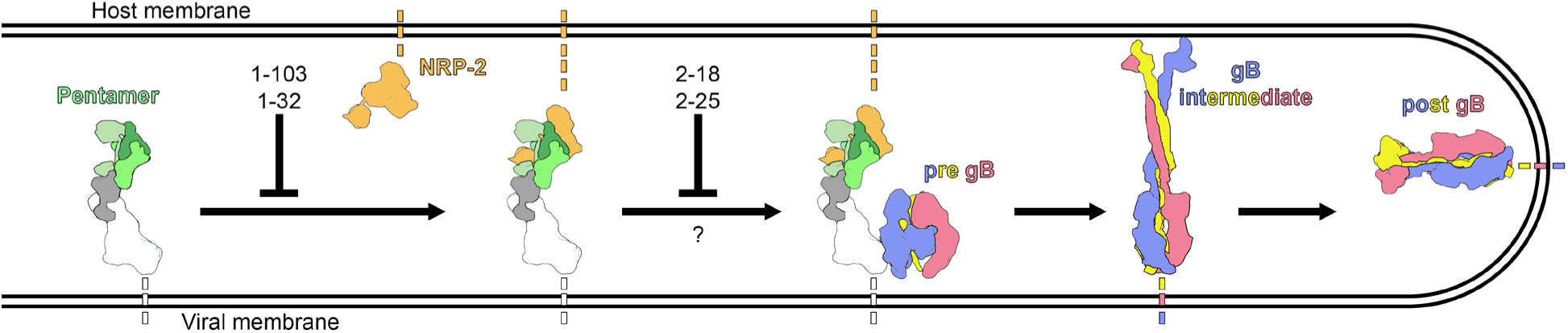
Pentamer-directed mAbs neutralize via distinct mechanisms. A cartoon is shown depicting the infection of an endothelial or epithelial cell by HCMV. Pentamer is colored according to Fig. 1, NRP2 is colored orange and the three protomers of gB are colored red, blue and yellow. The stages of infection that mAbs 1-103, 1-32, 2-18 and 2-25 are predicted to disrupt are denoted by inhibition arrows.

**Supplementary Table 1:** X-ray crystallographic data collection and refinement.

|  | 1-103 Fab | 1-32 Fab | 2-18 Fab | 2-25 Fab |
| --- | --- | --- | --- | --- |
| **Data collection** |  |  |  |  |
| Facility | APS 19-ID | APS 19-ID | APS 19-ID | APS 19-ID |
| Wavelength (Å) | 0.979 | 0.979 | 0.979 | 0.979 |
| Space group | *P*2 | *P*4<sub>1</sub> 2<sub>1</sub> 2 | *C*2 | *P*4<sub>3</sub> 2<sub>1</sub> 2 |
| Cell dimensions |  |  |  |  |
| a, b, c (Å) | 95.4, 105.2, 102.7 | 65.7, 65.7, 191.9 | 129.5, 59.2, 141.7 | 180.9, 180.9, 138.9 |
| $\alpha$, $\beta$, $\gamma$ (°) | 90.0, 91.4, 90.0 | 90.0, 90.0, 90.0 | 90.0, 91.5, 90.0 | 90.0, 90.0, 90.0 |
| Resolution range (Å) | 52.62-1.90 (1.93-1.90) | 54.20-2.10 (2.16-2.10) | 65.02-2.80 (2.95-2.80) | 69.89-2.51 (2.57-2.51) |
| R<sub>merge</sub> | 0.059 (0.407) | 0.174 (0.951) | 0.352 (1.283) | 0.187 (2.033) |
| CC1/2 | 0.996 (0.736) | 0.987 (0.753) | 0.851 (0.291) | 0.991 (0.522) |
| I/σI | 5.9 (1.8) | 6.0 (1.5) | 4.0 (1.7) | 9.3 (2.0) |
| Completeness (%) | 90.2 (91.3) | 99.9 (99.8) | 98.8 (96.7) | 98.6 (99.9) |
| Redundancy | 2.0 (2.0) | 7.2 (6.6) | 4.1 (3.8) | 6.8 (7.1) |
| **Refinement** |  |  |  |  |
| No. reflections | 143,434 (14,383) | 25,398 (2,476) | 26,469 (2,446) | 77,206 (7,680) |
| R<sub>work</sub>/R<sub>free</sub> (%) | 19.1/21.7 | 26.8/29.7 | 22.1/25.5 | 18.5/21.9 |
| No. non-hydrogen atoms |  |  |  |  |
| Protein | 12,781 | 3,226 | 6,726 | 9,633 |
| Ligand/ion | 0 | 0 | 0 | 0 |
| Water | 1,627 | 70 | 70 | 507 |
| B-factors (Å<sup>2</sup>) |  |  |  |  |
| Protein | 28.7 | 62.9 | 27.0 | 39.9 |
| Solvent | 38.8 | 49.7 | 27.1 | 42.9 |
| R.m.s. deviations |  |  |  |  |
| Bond lengths (Å) | 0.007 | 0.013 | 0.011 | 0.011 |
| Bond angles (°) | 1.33 | 1.41 | 1.46 | 1.23 |
| Ramachandran (%) |  |  |  |  |
| Favored | 98.4 | 95.7 | 97.3 | 98.1 |
| Allowed | 1.6 | 4.3 | 2.7 | 1.9 |
| Outliers | 0 | 0 | 0 | 0 |
| **PDB ID** | 7LYV | 7M1C | 7KBA | 7LYW |

**Supplementary Table 2:**
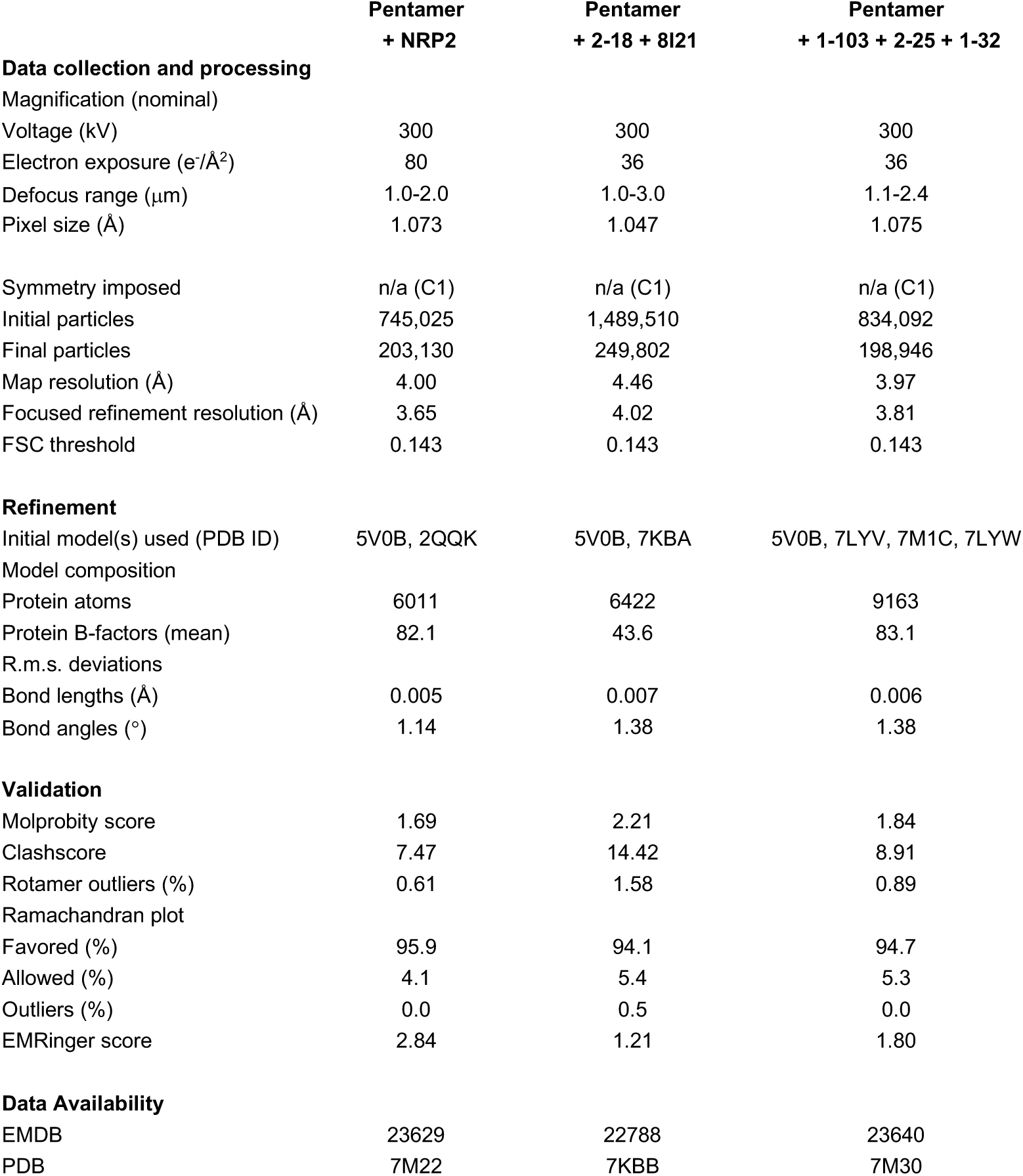
Cryo-EM data collection and refinement.

## REFERENCES

1 Gerna, G. et al. Dendritic-cell infection by human cytomegalovirus is restricted to strains carrying functional UL131-128 genes and mediates efficient viral antigen presentation to CD8+ T cells. J Gen Virol 86, 275–284, doi:10.1099/vir.0.80474-0 (2005).

2 Vanarsdall, A. L., Chase, M. C. & Johnson, D. C. Human cytomegalovirus glycoprotein gO complexes with gH/gL, promoting interference with viral entry into human fibroblasts but not entry into epithelial cells. J Virol 85, 11638–11645, doi:10.1128/JVI.05659-11 (2011).

3 Adler, B. et al. Role of human cytomegalovirus UL131A in cell type-specific virus entry and release. J Gen Virol 87, 2451–2460, doi:10.1099/vir.0.81921-0 (2006).

4 Martinez-Martin, N. et al. An Unbiased Screen for Human Cytomegalovirus Identifies Neuropilin-2 as a Central Viral Receptor. Cell 174, 1158–1171 e1119, doi:10.1016/j.cell.2018.06.028 (2018).

5 Teesalu, T., Sugahara, K. N., Kotamraju, V. R. & Ruoslahti, E. C-end rule peptides mediate neuropilin-1-dependent cell, vascular, and tissue penetration. Proc Natl Acad Sci U S A 106, 16157–16162, doi:10.1073/pnas.0908201106 (2009).

6 Tsai, Y. C. et al. Structural studies of neuropilin-2 reveal a zinc ion binding site remote from the vascular endothelial growth factor binding pocket. FEBS J 283, 1921–1934, doi:10.1111/febs.13711 (2016).

7 Cannon, M. J., Schmid, D. S. & Hyde, T. B. Review of cytomegalovirus seroprevalence and demographic characteristics associated with infection. Rev Med Virol 20, 202–213, doi:10.1002/rmv.655 (2010).

8 Ahlfors, K., Forsgren, M., Ivarsson, S. A., Harris, S. & Svanberg, L. Congenital cytomegalovirus infection: on the relation between type and time of maternal infection and infant’s symptoms. Scand J Infect Dis 15, 129–138, doi:10.3109/inf.1983.15.issue-2.01 (1983).

9 Pass, R. F., Fowler, K. B., Boppana, S. B., Britt, W. J. & Stagno, S. Congenital cytomegalovirus infection following first trimester maternal infection: symptoms at birth and outcome. J Clin Virol 35, 216–220, doi:10.1016/j.jcv.2005.09.015 (2006).

10 Roark, H. K., Jenks, J. A., Permar, S. R. & Schleiss, M. R. Animal Models of Congenital Cytomegalovirus Transmission: Implications for Vaccine Development. J Infect Dis 221, S60–S73, doi:10.1093/infdis/jiz484 (2020).

11 Ligat, G., Cazal, R., Hantz, S. & Alain, S. The human cytomegalovirus terminase complex as an antiviral target: a close-up view. FEMS Microbiol Rev 42, 137–145, doi:10.1093/femsre/fuy004 (2018).

12 Mumtaz, K. et al. Universal prophylaxis or preemptive strategy for cytomegalovirus disease after liver transplantation: a systematic review and meta-analysis. Am J Transplant 15, 472–481, doi:10.1111/ajt.13044 (2015).

13 McGeoch, D. J., Cook, S., Dolan, A., Jamieson, F. E. & Telford, E. A. Molecular phylogeny and evolutionary timescale for the family of mammalian herpesviruses. J Mol Biol 247, 443–458, doi:10.1006/jmbi.1995.0152 (1995).

14 Isaacson, M. K. & Compton, T. Human cytomegalovirus glycoprotein B is required for virus entry and cell-to-cell spread but not for virion attachment, assembly, or egress. J Virol 83, 3891–3903, doi:10.1128/JVI.01251-08 (2009).

15 Wang, D. & Shenk, T. Human cytomegalovirus virion protein complex required for epithelial and endothelial cell tropism. Proc Natl Acad Sci U S A 102, 18153–18158, doi:10.1073/pnas.0509201102 (2005).

16 Zhou, M., Lanchy, J. M. & Ryckman, B. J. Human Cytomegalovirus gH/gL/gO Promotes the Fusion Step of Entry into All Cell Types, whereas gH/gL/UL128-131 Broadens Virus Tropism through a Distinct Mechanism. J Virol 89, 8999–9009, doi:10.1128/JVI.01325-15 (2015).

17 Ciferri, C. et al. Structural and biochemical studies of HCMV gH/gL/gO and Pentamer reveal mutually exclusive cell entry complexes. Proc Natl Acad Sci U S A 112, 1767–1772, doi:10.1073/pnas.1424818112 (2015).

18 Kabanova, A. et al. Platelet-derived growth factor-alpha receptor is the cellular receptor for human cytomegalovirus gHgLgO trimer. Nat Microbiol 1, 16082, doi:10.1038/nmicrobiol.2016.82 (2016).

19 Wu, Y. et al. Human cytomegalovirus glycoprotein complex gH/gL/gO uses PDGFR-alpha as a key for entry. PLoS Pathog 13, e1006281, doi:10.1371/journal.ppat.1006281 (2017).

20 Kschonsak, M. et al. Structures of HCMV Trimer reveal the basis for receptor recognition and cell entry. Cell, doi:10.1016/j.cell.2021.01.036 (2021).

21 Hahn, G. et al. Human cytomegalovirus UL131-128 genes are indispensable for virus growth in endothelial cells and virus transfer to leukocytes. J Virol 78, 10023–10033, doi:10.1128/JVI.78.18.10023-10033.2004 (2004).

22 Ryckman, B. J. et al. Characterization of the human cytomegalovirus gH/gL/UL128-131 complex that mediates entry into epithelial and endothelial cells. J Virol 82, 60–70, doi:10.1128/JVI.01910-07 (2008).

23 Wille, P. T., Wisner, T. W., Ryckman, B. & Johnson, D. C. Human cytomegalovirus (HCMV) glycoprotein gB promotes virus entry in trans acting as the viral fusion protein rather than as a receptor-binding protein. mBio 4, e00332–00313, doi:10.1128/mBio.00332-13 (2013).

24 Soker, S., Takashima, S., Miao, H. Q., Neufeld, G. & Klagsbrun, M. Neuropilin-1 is expressed by endothelial and tumor cells as an isoform-specific receptor for vascular endothelial growth factor. Cell 92, 735–745, doi:10.1016/s0092-8674(00)81402-6 (1998).

25 Takagi, S., Tsuji, T., Amagai, T., Takamatsu, T. & Fujisawa, H. Specific cell surface labels in the visual centers of Xenopus laevis tadpole identified using monoclonal antibodies. Dev Biol 122, 90–100, doi:10.1016/0012-1606(87)90335-6 (1987).

26 Dallas, N. A. et al. Neuropilin-2-mediated tumor growth and angiogenesis in pancreatic adenocarcinoma. Clin Cancer Res 14, 8052–8060, doi:10.1158/1078-0432.CCR-08-1520 (2008).

27 Kitsukawa, T., Shimono, A., Kawakami, A., Kondoh, H. & Fujisawa, H. Overexpression of a membrane protein, neuropilin, in chimeric mice causes anomalies in the cardiovascular system, nervous system and limbs. Development 121, 4309–4318 (1995).

28 Prahst, C. et al. Neuropilin-1-VEGFR-2 complexing requires the PDZ-binding domain of neuropilin-1. J Biol Chem 283, 25110–25114, doi:10.1074/jbc.C800137200 (2008).

29 Takagi, S. et al. The A5 antigen, a candidate for the neuronal recognition molecule, has homologies to complement components and coagulation factors. Neuron 7, 295–307, doi:10.1016/0896-6273(91)90268-5 (1991).

30 Favier, B. et al. Neuropilin-2 interacts with VEGFR-2 and VEGFR-3 and promotes human endothelial cell survival and migration. Blood 108, 1243–1250, doi:10.1182/blood-2005-11-4447 (2006).

31 Parker, M. W., Xu, P., Li, X. & Vander Kooi, C. W. Structural basis for selective vascular endothelial growth factor-A (VEGF-A) binding to neuropilin-1. J Biol Chem 287, 11082–11089, doi:10.1074/jbc.M111.331140 (2012).

32 Daly, J. L. et al. Neuropilin-1 is a host factor for SARS-CoV-2 infection. Science 370, 861–865, doi:10.1126/science.abd3072 (2020).

33 E, X. et al. OR14I1 is a receptor for the human cytomegalovirus pentameric complex and defines viral epithelial cell tropism. Proc Natl Acad Sci U S A 116, 7043–7052, doi:10.1073/pnas.1814850116 (2019).

34 Vanarsdall, A. L. et al. CD147 Promotes Entry of Pentamer-Expressing Human Cytomegalovirus into Epithelial and Endothelial Cells. mBio 9, doi:10.1128/mBio.00781-18 (2018).

35 Ha, S. et al. Neutralization of Diverse Human Cytomegalovirus Strains Conferred by Antibodies Targeting Viral gH/gL/pUL128-131 Pentameric Complex. J Virol 91, doi:10.1128/JVI.02033-16 (2017).

36 Vanarsdall, A. L. et al. HCMV trimer- and pentamer-specific antibodies synergize for virus neutralization but do not correlate with congenital transmission. Proc Natl Acad Sci U S A 116, 3728–3733, doi:10.1073/pnas.1814835116 (2019).

37 Freed, D. C. et al. Pentameric complex of viral glycoprotein H is the primary target for potent neutralization by a human cytomegalovirus vaccine. Proc Natl Acad Sci U S A 110, E4997–5005, doi:10.1073/pnas.1316517110 (2013).

38 Appleton, B. A. et al. Structural studies of neuropilin/antibody complexes provide insights into semaphorin and VEGF binding. EMBO J 26, 4902–4912, doi:10.1038/sj.emboj.7601906 (2007).

39 Cohen-Dvashi, H., Kilimnik, I. & Diskin, R. Structural basis for receptor recognition by Lujo virus. Nat Microbiol 3, 1153–1160, doi:10.1038/s41564-018-0224-5 (2018).

40 Parker, M. W. et al. Structural basis for VEGF-C binding to neuropilin-2 and sequestration by a soluble splice form. Structure 23, 677–687, doi:10.1016/j.str.2015.01.018 (2015).

41 Vander Kooi, C. W., et al. Structural basis for ligand and heparin binding to neuropilin B domains. Proc Natl Acad Sci U S A 104, 6152–6157, doi:10.1073/pnas.0700043104 (2007).

42 Chandramouli, S. et al. Structural basis for potent antibody-mediated neutralization of human cytomegalovirus. Sci Immunol 2, doi:10.1126/sciimmunol.aan1457 (2017).

43 Xia, L. et al. Active evolution of memory B-cells specific to viral gH/gL/pUL128/130/131 pentameric complex in healthy subjects with silent human cytomegalovirus infection. Oncotarget 8, 73654–73669, doi:10.18632/oncotarget.18359 (2017).

44 Ye, X. et al. Recognition of a highly conserved glycoprotein B epitope by a bivalent antibody neutralizing HCMV at a post-attachment step. PLoS Pathog 16, e1008736, doi:10.1371/journal.ppat.1008736 (2020).

45 Yelland, T. & Djordjevic, S. Crystal Structure of the Neuropilin-1 MAM Domain: Completing the Neuropilin-1 Ectodomain Picture. Structure 24, 2008–2015, doi:10.1016/j.str.2016.08.017 (2016).

46 Sinzger, C. et al. Fibroblasts, epithelial cells, endothelial cells and smooth muscle cells are major targets of human cytomegalovirus infection in lung and gastrointestinal tissues. J Gen Virol 76 (Pt 4), 741–750, doi:10.1099/0022-1317-76-4-741 (1995).

47 Si, Z. et al. Different functional states of fusion protein gB revealed on human cytomegalovirus by cryo electron tomography with Volta phase plate. PLoS Pathog 14, e1007452, doi:10.1371/journal.ppat.1007452 (2018).

48 Ciferri, C. et al. Antigenic Characterization of the HCMV gH/gL/gO and Pentamer Cell Entry Complexes Reveals Binding Sites for Potently Neutralizing Human Antibodies. PLoS Pathog 11, e1005230, doi:10.1371/journal.ppat.1005230 (2015).

49 Gerna, G., Percivalle, E., Perez, L., Lanzavecchia, A. & Lilleri, D. Monoclonal Antibodies to Different Components of the Human Cytomegalovirus (HCMV) Pentamer gH/gL/pUL128L and Trimer gH/gL/gO as well as Antibodies Elicited during Primary HCMV Infection Prevent Epithelial Cell Syncytium Formation. J Virol 90, 6216–6223, doi:10.1128/JVI.00121-16 (2016).

50 Su, H. et al. Potent Bispecific Neutralizing Antibody Targeting Glycoprotein B and the gH/gL/pUL128/130/131 Complex of Human Cytomegalovirus. Antimicrob Agents Chemother 65, doi:10.1128/AAC.02422-20 (2021).

51 Jones, H. G. et al. Iterative screen optimization maximizes the efficiency of macromolecular crystallization. Acta Crystallogr F Struct Biol Commun 75, 123–131, doi:10.1107/S2053230X18017338 (2019).

52 Battye, T. G., Kontogiannis, L., Johnson, O., Powell, H. R. & Leslie, A. G. iMOSFLM: a new graphical interface for diffraction-image processing with MOSFLM. Acta Crystallogr D Biol Crystallogr 67, 271–281, doi:10.1107/S0907444910048675 (2011).

53 Evans, P. R. & Murshudov, G. N. How good are my data and what is the resolution? Acta Crystallogr D Biol Crystallogr 69, 1204–1214, doi:10.1107/S0907444913000061 (2013).

54 McCoy, A. J. Solving structures of protein complexes by molecular replacement with Phaser. Acta Crystallogr D Biol Crystallogr 63, 32–41, doi:10.1107/S0907444906045975 (2007).

55 Emsley, P. & Cowtan, K. Coot: model-building tools for molecular graphics. Acta Crystallogr D Biol Crystallogr 60, 2126–2132, doi:10.1107/S0907444904019158 (2004).

56 Adams, P. D. et al. PHENIX: building new software for automated crystallographic structure determination. Acta Crystallogr D Biol Crystallogr 58, 1948–1954, doi:10.1107/s0907444902016657 (2002).

57 Croll, T. I. ISOLDE: a physically realistic environment for model building into low-resolution electron-density maps. Acta Crystallogr D Struct Biol 74, 519–530, doi:10.1107/S2059798318002425 (2018).

58 Morin, A. et al. Collaboration gets the most out of software. Elife 2, e01456, doi:10.7554/eLife.01456 (2013).

59 Mastronarde, D. N. SerialEM: A Program for Automated Tilt Series Acquisition on Tecnai Microscopes Using Prediction of Specimen Position. Microscopy and Microanalysis, doi:10.1017/S1431927603445911 (2003).

60 Carragher, B. et al. Leginon: an automated system for acquisition of images from vitreous ice specimens. J Struct Biol 132, 33–45, doi:10.1006/jsbi.2000.4314 (2000).

61 Tegunov, D. & Cramer, P. Real-time cryo-electron microscopy data preprocessing with Warp. Nat Methods 16, 1146–1152, doi:10.1038/s41592-019-0580-y (2019).

62 Punjani, A., Rubinstein, J. L., Fleet, D. J. & Brubaker, M. A. cryoSPARC: algorithms for rapid unsupervised cryo-EM structure determination. Nat Methods 14, 290–296, doi:10.1038/nmeth.4169 (2017).

63 Punjani, A., Zhang, H. & Fleet, D. J. Non-uniform refinement: adaptive regularization improves single-particle cryo-EM reconstruction. Nat Methods 17, 1214–1221, doi:10.1038/s41592-020-00990-8 (2020).

64 Sanchez-Garcia, R. et al. DeepEMhancer: a deep learning solution for cryo-EM volume post-processing. bioRxiv, doi:https://doi.org/10.1101/2020.06.12.148296 (2020).

65 Pettersen, E. F. et al. UCSF Chimera--a visualization system for exploratory research and analysis. J Comput Chem 25, 1605–1612, doi:10.1002/jcc.20084 (2004).

66 Li, L. et al. Potent neutralizing antibodies elicited by dengue vaccine in rhesus macaque target diverse epitopes. PLoS pathogens 15, e1007716, doi:10.1371/journal.ppat.1007716 (2019).

67 Ye, X. et al. Structural Basis for Recognition of Human Enterovirus 71 by a Bivalent Broadly Neutralizing Monoclonal Antibody. PLoS pathogens 12, e1005454, doi:10.1371/journal.ppat.1005454 (2016).

68 Tang, A. et al. A novel high-throughput neutralization assay for supporting clinical evaluations of human cytomegalovirus vaccines. Vaccine 29, 8350–8356, doi:10.1016/j.vaccine.2011.08.086 (2011).

69 Ye, X. et al. Identification of adipocyte plasma membrane-associated protein as a novel modulator of human cytomegalovirus infection. PLoS pathogens 15, e1007914, doi:10.1371/journal.ppat.1007914 (2019).

